# Developmental system drift in dorsoventral patterning is linked to transitions to autonomous development in Annelida

**DOI:** 10.1101/2025.05.29.656861

**Authors:** Allan M. Carrillo-Baltodano, Emmanuel Haillot, Steffanie Mutiara Meha, Imran Luqman, Artenis Pashaj, Yun-Ju Lee, Tsai-Ming Lu, David E. K. Ferrier, Stephan Q. Schneider, José M. Martín-Durán

## Abstract

The Bone Morphogenetic Protein (BMP) pathway is the ancestral signalling system defining the dorsoventral axis in bilaterally symmetrical animals. However, Spiralia, a large bilaterian clade including molluscs and annelids, uses the Fibroblast Growth Factor pathway and ERK1/2 as the ancestral cue to establish their posterodorsal side. How this profound change in axial patterning evolved and what it implied for BMP’s developmental role remains elusive. Here, we studied four annelid species and combined disruption of the BMP and Activin/Nodal pathways with transcriptomics and blastomere deletions to demonstrate that BMP is ancestrally downstream of ERK1/2 and promotes dorsoventral development in Spiralia. Importantly, this signalling hierarchy is lost in annelids that secondarily transitioned into a maternally controlled, autonomous development. While some, like *Capitella teleta*, use Activin/Nodal, *Platynereis dumerilii* relies on BMP to establish dorsoventral polarity only in the head.

Unexpectedly, this divergence in upstream axial regulators implied extensive rewiring of downstream targets, as inferred by comparing *C. teleta* and *Owenia fusiformis*. Our data clarify the ancestral axial role for BMP in Spiralia, unveiling a potential causal link between parallel shifts to autonomous cell-fate specification in early development and the emergence of developmental system drift, a pervasive yet poorly understood phenomenon in animal embryogenesis.

## Main text

More than 99% of the known animal species exhibit bilateral symmetry^1,2^, with their anatomies organised along two orthogonal symmetry planes referred to as the primary anteroposterior (AP) and the secondary dorsoventral (DV) axes. Remarkably, distantly related bilaterian lineages use homologous developmental signalling pathways to specify their body axes^3,4^. In Deuterostomia (i.e., chordates, echinoderms, and hemichordates) and Ecdysozoa (e.g., arthropods), the Bone Morphogenetic Protein (BMP) pathway specifies the DV axis through the interplay of extracellular secreted ligands (BMP2/4) and antagonists (Chordin). This creates an intracellular morphogenetic gradient of active phosphorylated SMAD1/5/8 (pSMAD1/5/8) that accumulates in the opposite pole to the source of the antagonist, promoting dorsal structures in non-chordates and ventral in chordates^5^ (Fig. 1a). However, in Spiralia (e.g. molluscs and annelids), the third major lineage of bilaterian animals^6^ characterised by an ancestral stereotypical early embryonic program named spiral cleavage, the initial specification of the DV axis ancestrally involved the Fibroblast Growth Factor receptor (FGFR) and ERK1/2 pathways^7–9^ (Fig. 1a). Notably, an opposing gradient along the secondary body axis of BMP antagonists and pSMAD1/5/8 also occurs in Cnidaria^10^, the sister group to Bilateria^2^. Therefore, an upstream role of the BMP pathway in defining the body symmetry and DV axis is likely ancestral to Bilateria^2,5^ (Fig. 1a), rendering the condition in Spiralia a pronounced and unique yet still poorly understood transition in the mechanisms controlling animal body patterning.

**Figure 1.**
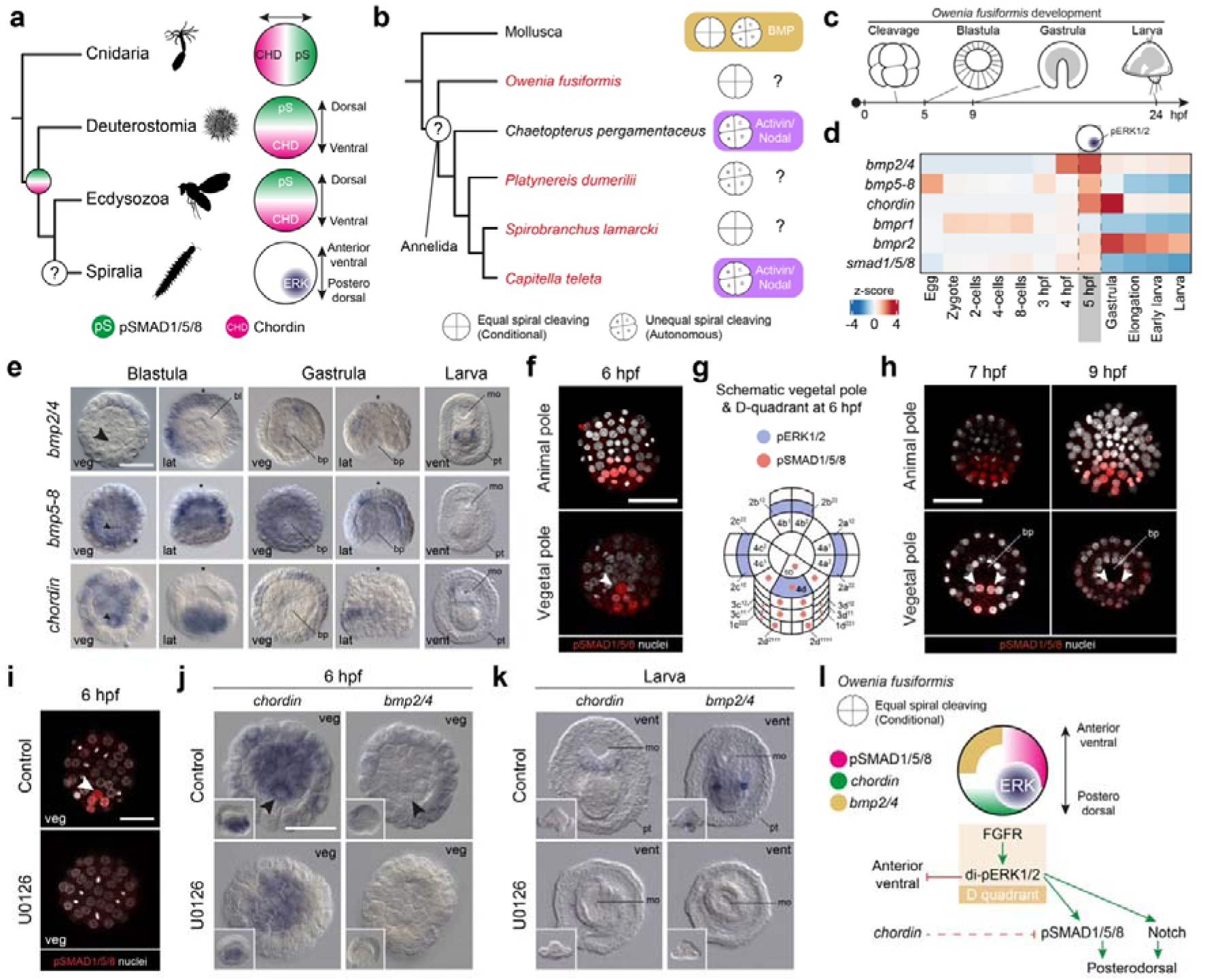
A dorsoventral gradient of pSMAD1/5/8 in the annelid *Owenia fusiformis*. (**a**) BMP and BMP antagonists (e.g. Chordin) set up a gradient of pSMAD1/5/8 that defines the secondary axis in cnidarians and the dorsoventral (DV) axis in bilaterians. In Spiralia, one cell activated by ERK acts as an axial organiser, and BMP’s role in DV patterning is contentious. (**b**) In the studied annelid species, *C. teleta* and *C. pergamentaceus*, which have autonomous/unequal spiral cleavage, Activin/Nodal specifies the DV axis. Whether this represents the ancestral annelid state is unclear. To solve this, we have studied four annelids (in red) with different modes of spiral cleavage. (**c**) Schematic diagram of O. fusiformis’ embryogenesis. (**d**) Heatmap of the relative expression of members of the BMP pathway during the development of *O. fusiformis.* (**e**) Whole mount *in situ* hybridisation of a subset (ligands and antagonist) of the BMP pathway genes in blastula (6 hours post-fertilisation, hpf), gastrula (9 hpf) and larval stages (24 hpf). (**f**, **h, i**) Z-stack projections of embryos at 6-, 7-, and 9 hpf stained against pSMAD1/5/8 (red) and nuclei (DAPI). (**g**) Schematic representation of the vegetal pole during the activation of pSMAD1/5/8. Cell’s identity follows the standard spiralian nomenclature. (**j**, **k**) Whole mount in situ hybridisation of *chordin* and *bmp2/4* demonstrates that inhibition of ERK1/2 activation prevents BMP activation. (**l**) Schematic representation of the signalling regulatory interactions specifying the bilateral symmetry in *O. fusiformis*. Arrowheads in **e**, **f** and **h**, **i** point to the embryonic organiser cell (4d) or its subsequent daughter cells 4d^1^ and 4d^2^. An asterisk marks the animal/apical. Scale bars are 50 µm. an anus, bl blastocoel, bp blastopore, lat lateral, mo mouth, pt prototroch, veg vegetal, vent ventral. Drawings are not to scale.

Concomitant with the evolution of FGFR-ERK1/2-mediated axial patterning, the developmental role of the BMP signalling pathway diversified within Spiralia^5,11^. This is particularly evident in molluscs and annelids, where an autonomous, maternally controlled mode of specifying the initial embryonic axes evolved independently multiple times from the ancestral FGFR-ERK1/2-mediated conditional spiral cleavage^7,11^. While in some molluscs, BMP is downstream of ERK1/2 activity and controls DV development ––whilst in some cases promoting and in others repressing neurogenesis^9,12^–– BMP controls only head and anterior neural development in others^13^. Surprisingly, BMP does not seem to play a role in DV specification in annelids^14–17^. However, only species with the derived autonomous mode of spiral cleavage have been studied (Fig. 1b). Moreover, a complex pattern of gene loss characterises the evolutionary history of *chordin* in annelids^18^. Accordingly, related signalling pathways, such as Activin/Nodal, and lineage-specific signalling interactions specify the annelid DV axis^14–16,19^. To complicate things further, brachiopods, a spiralian lineage that has lost the ancestral spiral cleavage^20^, exhibit a Deuterostomia- and Ecdysozoa-like condition, where BMP signalling controls the DV embryonic polarity, and FGFR participates in late stages of mesoderm development^21–23^. Therefore, the ancestral role of the BMP signalling pathway remains contentious in Spiralia, preventing a fundamental understanding of how the shift to an FGFR-ERK1/2-mediated DV axial cue originated during animal evolution.

Here, we apply a phylum-wide comparative approach to resolve the role of the BMP and Activin/Nodal signalling pathways in Annelida. By studying two distantly related annelids with the ancestral conditional spiral cleavage, *Owenia fusiformis* and *Spirobranchus lamarcki*, we demonstrate that ERK1/2 activates BMP signalling on the prospective dorsal side of the embryo, which is required to activate downstream genes involved in the development of dorsoposterior structures. This is like what has been described in a mollusc with conditional spiral cleavage^12,24^ and likely represents the ancestral condition for Annelida and Spiralia. However, the annelid *Platynereis dumerilii*, a species with the derived autonomous spiral cleavage, restricts the axial patterning role of the BMP pathway to the head region. Notably, this is unlike other annelids with autonomous development, such as *Capitella teleta* and *Chaetopterus pergamentaceus*, which use Activin/Nodal instead^15,16,19^. In *P. dumerilii*, Activin/Nodal does not affect DV polarity. Still, it contributes to the normal development of DV structures in *O. fusiformis*, albeit through regulating different downstream genes than the BMP signalling. Altogether, our findings resolve the ancestral axial role of the BMP pathway in Spiralia, further supporting a deep evolutionary conservation of the body patterning mechanisms across spiralians with the ancestral conditional mode of cell fate specification. Importantly, parallel independent transitions into autonomous development co-occurred with developmental system drift (DSD) in DV patterning in Annelida, uncovering a unique and tractable system to explore the evolutionary forces and developmental mechanisms underpinning DSD, a widespread yet poorly understood phenomenon in evolutionary developmental biology.

## Results

### A dorsoventral gradient of pSMAD1/5/8 in O. fusiformis

To elucidate the role of BMP signalling in *O. fusiformis*, an early-branching annelid that develops via the ancestral state of conditional spiral cleavage, we first looked at the expression dynamics of the core components of this pathway during embryogenesis (Fig. 1c– e). The expression of the ligands *bmp2/4* and *bmp5-8* peaks at 5 hours post fertilisation (hpf) when the axial organising cell ––the 4d blastomere–– gets specified (Fig. 1d). At 5 and 6 hpf, *bmp2/4* is expressed on the anterior region of the embryo, overlapping anteriorly with the broad vegetal expression of the antagonist *chordin*^18^, whose expression peaks with gastrulation (Fig. 1d, e). At that stage (9 hpf), *bmp2/4*, *bmp5-8*, and *chordin* are restricted to non-overlapping, staggered expression domains in the presumptive anteroventral ectoderm of the embryo (based on the position of 4d and its progeny), with *chordin* expressed more vegetally and *bmp5-8* closer to the animal pole (Fig. 1e). However, by the larval stage (24 hpf), *chordin* and *bmp2/4* are expressed in separate areas, surrounding the mouth and just beneath the hindgut, respectively (Fig. 1e). Unlike the extracellular signalling players, the two BMP receptors^25^, *bmpr1* and *bmpr2*, and the secondary messenger *smad1/5/8* are ubiquitously expressed in the blastula and gastrula stages (Extended Data Fig. 1a), suggesting that most, if not all, cells could respond to BMP signalling during axial patterning despite the localised expression of the ligands and antagonists in *O. fusiformis*.

To identify the region with active BMP signalling during axial patterning in *O. fusiformis*, we next localised the activation of pSMAD1/5/8 with a cross-reactive antibody (Fig. 1f–h; Extended Data Fig. 1d). pSMAD1/5/8 is not detected at 5 hpf during the specification of the embryonic organiser^7^, but immediately after, at 6 hpf, in the organiser and cells of the dorsoposterior region extending from the animal to the vegetal pole (Fig. 1f, g). This pattern remains as gastrulation proceeds, with pSMAD1/5/8 remaining enriched in the progeny of the embryonic organiser and the prospective dorsoposterior half of the embryo (Fig. 1h).

Notably, active pSMAD1/5/8 occurs on the opposite side to the area of expression of *bmp2/4*, *bmp5-8*, and *chordin* (Fig. 1e). Thus, there is a gradient of pSMAD1/5/8 activity along the DV axis in the embryos of *O. fusiformis,* which, as in other bilaterians and cnidarians^5,26^, might emerge from the shuttling of BMP ligands by antagonists from the ventral ––where the two signalling molecules are co-expressed–– to the dorsal side.

Given that the first signs of pSMAD1/5/8 activity occur after the FGFR-ERK1/2-mediated signalling event defining the bilateral symmetry of the embryo^7^, we hypothesised that ERK1/2 activity could control the activation of pSMAD1/5/8 on the presumptive dorsoposterior side. To test this, we explored the protein domain composition of pSMAD1/5/8 to scan for ERK1/2-mediated phosphorylation sites and treated embryos with U0126, a specific inhibitor of ERK1/2 activation that prevents the specification of the axial embryonic organiser^7^. In vertebrates, ERK regulates SMAD1 via phosphorylation of residues in a linker region between two MAD domains^27–29^. However, these residues are only conserved across vertebrates and in invertebrate deuterostomes to a certain extent (Extended Data Fig. 1b, c). Treatment with U0126 from 4 to 6 hpf completely blocked the activation of pSMAD1/5/8 (Fig. 1i; Supplementary Table 1), downregulating the expression of *bmp2/4* and *chordin* at the gastrula and larval stages, which show a radialised phenotype (Fig. 1j, k; Supplementary Table 2). Therefore, FGFR-ERK1/2 establishes the bilateral symmetry in *O. fusiformis* embryos by regulating BMP signalling and the activation of pSMAD1/5/8 in the prospective dorsoposterior side, as also observed in some molluscs^9,12,24^ (Fig. 1l).

### BMP signalling specifies the DV axis in O. fusiformis

To investigate the role of the pSMAD1/5/8 DV gradient in axial patterning, we altered the BMP pathway by exposing *O. fusiformis* embryos with the specific BMP receptor inhibitor dorsomorphin homologue 1 (DMH1) and recombinant zebrafish BMP4 (rBMP4) from 4 to 6 hpf (Fig. 2a, b). Compared to the control condition, most (77%) larvae of DMH1-exposed embryos lack the chaetal sac but have a well-formed gut (Fig. 2c, d; Supplementary Table 3– 4). Conversely, most (86%) larvae of embryos treated with rBMP4 have a radial-like morphology, with only one gut opening surrounded by multiple ectopic chaetal sacs (Fig. 2c, d; Supplementary Table 3–4). Both treatments dysregulated pSMAD1/5/8 levels but not the specification of the embryonic organiser (as observed by the presence of the large 4d cell), with DMH1 exposure leading to a complete inhibition of SMAD1/5/8 and rBMP4 activating pSMAD1/5/8 in all embryonic cells, as expected by the ubiquitous expression of the receptor (Fig. 2e; Supplementary Table 1). Consistent with the morphological phenotypes, DMH1 treatment resulted in the expansion of the oral ectodermal marker *gsc*^7,23^, the loss of the dorsal markers *notch-like* and *BAMBI*^7^ but not the expression of the hindgut marker *cdx*, indicating the larvae have an AP axis but lack DV polarity (Fig. 2f, g; Supplementary Table 5–6). However, rBMP treatment resulted in the loss of *gsc* expression (but not of *cdx*) and the expansion of *notch-like* and *BAMBI* (Fig. 2f–g; Supplementary Table 5–6). Therefore, BMP signalling establishes a gradient of pSMAD1/5/8 activity that is necessary and sufficient to specify the dorsoventral polarity in *O. fusiformis*.

**Figure 2.**
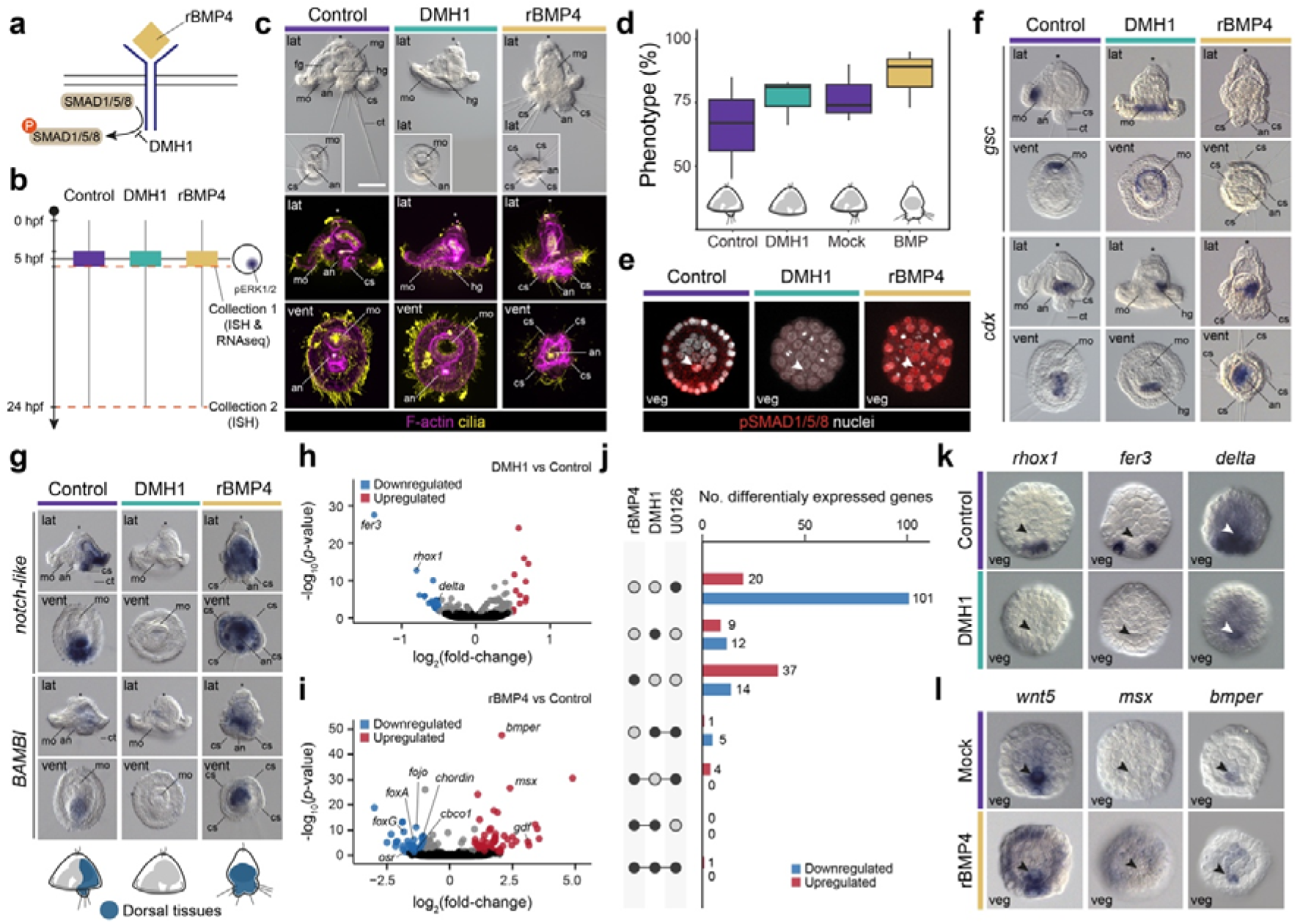
BMP controls a dorsoposterior gene regulatory network in *O. fusiformis*. (**a**) Simplified schematic of the BMP pathway and mode of action of DMH1 and recombinant BMP4 (rBMP4). (**b**) Schematic representation of the experimental design for the treatment windows and sample collections. (**c**) Differential interference contrast and z-stack projections of control and DMH1/rBMP4 larvae treated from 4 to 6 hpf. Insets in the first row are ventral views. Cilia (yellow) are labelled with tubulin, and F-actin (magenta) is labelled with phalloidin. (**d**) Percentage of phenotypes for each treatment. DMSO and mock control larvae have a wild-type morphology. DMH1 has a reduced dorsoposterior region, while rBMP4- treated larvae have ectopic dorsoposterior tissue. (**e**) Z-stack projections of blastula (6 hpf) treated with control and DMH1/rBMP4 and stained with pSMAD1/5/8. (**f, g**) Whole-mount *in situ* hybridisation of control and DMH1/rBMP4 larvae treated from 4 to 6 hpf for anterior (*gsc*), posterior (*cdx*) and the dorsal markers *notch-like* and *BAMBI*. (**h**, **i**) Volcano plots showing differentially expressed genes after DMH1 and rBMP4 treatment at the blastula stage (6 hpf), treated from 4 to 6 hpf. (**j**) Number of differentially expressed genes in the different treatments, compared with the changes in gene expression at 5.5 hpf observed after ERK1/2 inhibition. Full circles connected with a line indicated shared differentially expressed genes. (**k**, **l**) Validation via whole-mount *in situ* hybridisation of genes downregulated with DMH1 and upregulated with rBMP4 at the blastula stage (6 hpf). In all panels, the arrowheads point to the 4d organiser and asterisks mark the animal/apical pole. Scale bars are 50 µm. an anus, bp blastopore, ch chaetae, cs chaetal sac, fg foregut, lat lateral, mo mouth, pt prototroch, vent ventral, veg vegetal. Drawings are not to scale.

To dissect the gene regulatory network controlled by BMP signalling, we performed comparative bulk transcriptomic profiling between control, DMH1, and rBMP4-treated embryos at the onset of pSMAD1/5/8 activation at 6 hpf (Fig. 2b; Extended Data Fig. 2a, b). Differential expression analyses uncovered 12 upregulated and 19 downregulated genes after DMH1 treatment and 68 upregulated and 53 downregulated genes after rBMP4 (Fig. 2h, i; Supplementary Table 7–11). Notably, five of the nine downregulated genes after DMH1 treatment are also downregulated after inhibition of ERK1/2 di-phosphorylation^7^, including *rhox1*, *fer3*, and *delta* (Fig. 2j–k; Supplementary Table 12). Unlike the first two, whose expression entirely depends on normal BMP signalling, *delta* remains expressed in the embryonic organiser, suggesting that pSMAD1/5/8 activity mediates Notch/Delta signalling on the dorsal side but not in the 4d cell. Consistent with the morphological phenotype, rBMP4 treatment downregulates genes involved in anteroventral fates, such as *chordin*, *odd-skipped related* (*osr*) and *foxG* (Extended Data Fig. 2d, f) and expands genes localised in the embryonic organiser and other dorsoposterior blastomeres, such as *wnt5*, *msx*, and *bmper* (*crossveinless*) (Fig. 2l; Supplementary Table 13). Altogether, our data demonstrate that BMP signalling is required to activate dorsoposterior genes and sustain Notch/Delta activity while repressing anteroventral fates during axial patterning in *O. fusiformis*.

### BMP signalling does not affect neurogenesis in O. fusiformis

Associated with its role in setting up the DV axis, BMP signalling inhibits neural fates in vertebrates and arthropods^3,4^. Whether this also occurs in spiralians remains unclear^30^. In *O. fusiformis*, GO terms related to neural development are enriched after disturbing BMP signalling during axial patterning (Extended Data Fig. 2e, f). To further investigate the interplay between BMP signalling and neural development in this annelid, we analyse the expression of the pan-neural genes *elav1* and *synaptotagmin1* (*syt1*)^31,32^ and the conserved anterior neural marker *six3/6*, as well as the localisation of the neuropeptide RYamide^31,33–35^ (Extended Data Fig. 3). The nervous system of the *O. fusiformis* larva consists of a neuropeptide-rich apical organ connected bilaterally to a neurite ring that surrounds and presumably controls the movement of the primary ciliary band, the prototroch^31,32,36^ (Extended Data Fig. 3a). After DMH1 treatment (4 to 6 hpf), larvae lack expression of *syt1*, *elav1* and *six3/6* in the apical organ (Extended Data Fig. 3b–c; Supplementary Table 15).

However, there are RYamidergic neurons in the apical organ, although the nervous system around the prototroch is disorganised and less developed (Extended Data Fig. 3b; Supplementary Table 16). Paradoxically, rBMP4 treatment did not result in a lack of neural structures either, as expected if BMP signalling had an anti-neural role like in vertebrates and arthropods. Instead, *elav1*, *syt1* and *six3/6* are expressed in the apical pole of rBMP4-treated larvae, which exhibit fewer RYamidergic neurons (Extended Data Fig. 3c, e; Supplementary Tables 15–16). Therefore, even though it is difficult to discern whether the defects in the larval nervous system are a direct consequence of dysregulating BMP signalling or a result of the overall loss of axial identities after DMH1 and rBMP4 treatments, our data does not support a neural-related role coupled to the axial patterning function of the BMP signalling pathway in *O. fusiformis*.

### The role of ERK1/2 and BMP signalling in DV patterning is conserved in Annelida

The role of ERK1/2 and BMP signalling in setting the axial polarity in *O. fusiformis* markedly differs from the condition described in annelids with autonomous spiral cleavage, such as *C. pergamentaceus* and *C. teleta*, which use Activin/Nodal to set their DV axes^15,16,19^. To assess whether the axial patterning mechanisms found in *O. fusiformis* are conserved in other annelids with a similar conditional mode of spiral cleavage, we investigated the annelid *Spirobranchus lamarcki*^37^, a member of Serpulidae that is more closely related to *C. teleta* and *Helobdella robusta*. Similarly to *O. fusiformis* and another serpulid, *Hydroides exogonus*^38^, the embryos of *S. lamarcki* have a single cell enriched with dp-ERK1/2, which we deemed the 4d micromere (Fig. 3a). With organogenesis, these embryos develop into a typical trochophore larva with an anterior episphere (i.e., head) and a posterior hyposphere (i.e., trunk) separated by two ciliary bands, the prototroch and the metatroch^39^ (Fig. 3b). In these larvae, the anus is not terminal but opens slightly dorsally at the end of the hyposphere (Fig. 3b). Treatment with U0126 blocks the activation of ERK1/2, and also of SMAD1/5/8 (Fig. 3c; Supplementary Tables 17–18). However, the inhibition of the BMP signalling with DMH1 does not affect the dpERK1/2 activation and the specification of the 4d organiser (Fig. 3c; Supplementary Tables 17–18). In both cases, the inhibition of ERK1/2 and BMP signalling results in the loss of the hyposphere and the anus at 48 hpf, with DMH1-treated larvae exhibiting more marked anteroposterior elongation than those developing after inhibition of ERK1/2 (Fig. 3d). Conversely, treatment with rBMP4 leads to larvae with a reduced episphere, lacking a mouth and foregut muscles but presenting a dorsal anus (Fig. 3e). Therefore, ERK1/2 and BMP signalling establish the bilateral and DV polarity in *S. lamarcki*, respectively, demonstrating that distantly related annelids and molluscs with conditional spiral cleavage use homologous signalling interactions to define their axial patterning.

**Figure 3.**
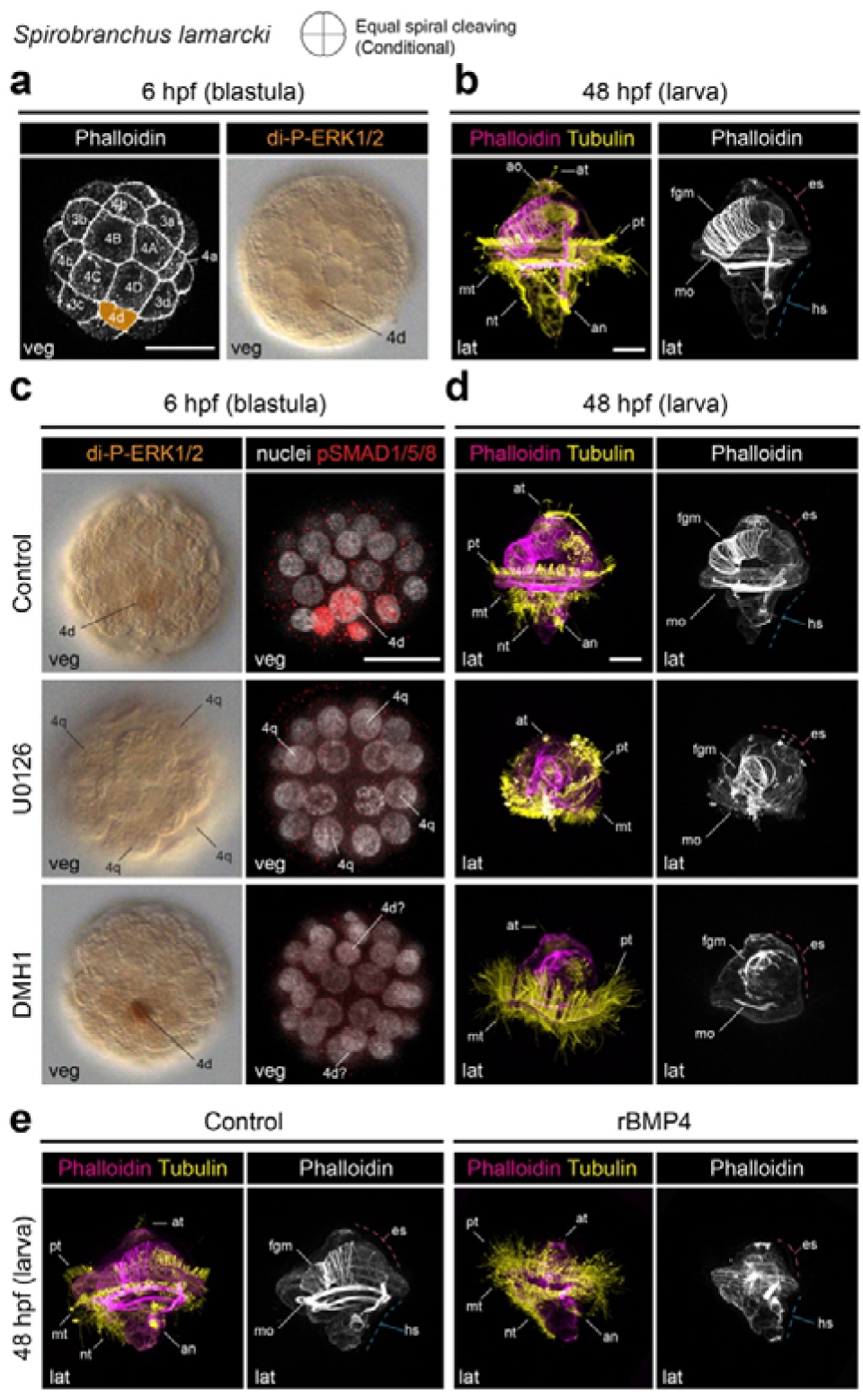
The role of ERK1/2 and BMP in the annelid with conditional cleavage *Spirobranchus lamarcki*. (**a**) Z-stack projections and Differential Interference Contrast (DIC) of blastula (6 hpf) with cell membranes labelled with phalloidin (left) and anti-dp- ERK1/2 (right), which localises in the putative 4d cell. (**b**) Z-stack projections of a 48 hpf larva. Cilia (yellow) are labelled with tubulin, and F-actin (magenta/white) is labelled with phalloidin. (**c**) DIC images of control and U0126/DMH1-treated blastula (6 hpf) stained with dp-ERK1/2 and z-stack projections of control and U0126/DMH1-treated blastula (6 hpf) stained with DAPI (white) and pSMAD1/5/8 (red). (**d**) Z-stack projections of 48 hpf larvae developed from control and U0126/DMH1-treated embryos. Cilia (yellow) are labelled with tubulin, and F-actin (magenta/white) is labelled with phalloidin. (**d**) Z-stack projections of 48 hpf larvae developed from control and rBMP4-treated embryos. Cilia (yellow) are labelled with tubulin, and F-actin (magenta/white) is labelled with phalloidin. Scale bars are 25 µm. an anus, ao apical organ, at apical tuft, es episphere, fgm foregut muscle, hs hyposphere, lat lateral, mo mouth, pt prototroch, mt metatroch, nt neurotroch, tt telotroch, veg vegetal.

### BMP signalling patterns the larval head along the DV axis in P. dumerilii

The ancestral role of BMP signalling in DV patterning suggests that using Activin/Nodal in DV specification evolved convergently in *C. teleta* and *C. pergamentaceus*. To assess whether this represents a recurring feature among other annelids with autonomous development, we investigated the role of BMP signalling in *Platynereis dumerilii*, a well-established model annelid belonging to Errantia^40^, the sister clade to the group comprising *S. lamarcki* and *C. teleta* (Fig. 1b). As in other spiralians exhibiting unequal, autonomous cleavage, the two larger animal micromeres at the 8-cell stage ––1c and 1d–– give rise to the dorsal head region, while the two smaller micromeres ––1a and 1b–– contribute to the ventral head region in *P. dumerilii*^41–43^ (Fig. 4a). In subsequent cell divisions, the largest macromere 1D generates two uniquely large blastomeres ––2d and 4d–– which demarcate the dorsal side of the trunk (Fig. 4a). Thus, in contrast to *O. fusiformis* and *S. lamarcki*, early unequal cell divisions in *P. dumerilli* predetermine DV patterning, which arises independently of ERK1/2 activity ^44^. Although BMP influences DV patterning in the 2-day-old larva in *P. dumerilii*^14^, it remains unclear whether this signalling is essential for DV polarity during embryogenesis.

**Figure 4.**
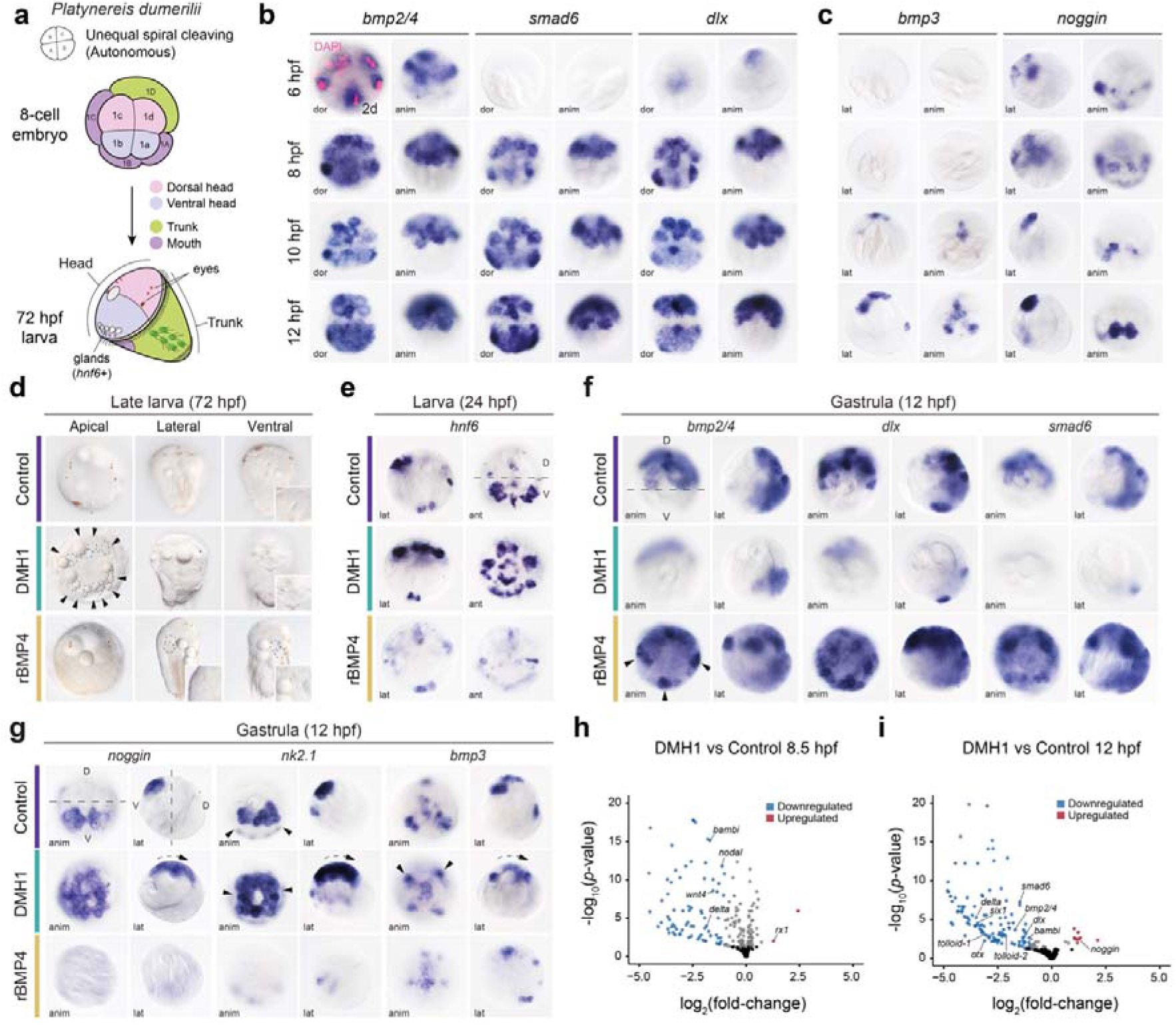
BMP controls the dorsoposterior axis of the head in *P. dumerilii*. (**a**) Diagram showing the origin of the dorsoventral structures from the 8 cell-stage embryo and their simplified fate map in the 72 hpf larva (see text for details). Three pairs of dorsal eyes and five ventral *hnf6*+ gland cells are shown. (**b**–**c**) Whole mount *in situ* hybridisation of the dorsoventral markers *bmp2/4*, *smad6*, *dlx*, *bmp3* and *noggin* at 6, 8, 10 and 12 hpf embryos. (**d)** Differential Interference Contrast (DIC) images of control, DMH1, and rBMP4-treated 72 hpf larvae. Arrowheads point to the radialisation of the gland cells (dashed) after DMH1 treatment. Dashed circles show ectopic mouths after rBMP4 treatment. (**e**–**g**) Whole mount *in situ* hybridisation of ventral (*hnf6*, *noggin*, *nk2.1* and *bmp3*) and dorsal (*bmp2/4*, *dlx* and *smad6*) markers in control, DMH1, and rBMP4-treated (**e)** 24 hpf larvae and (**f**–**g**) 12 hpf gastrulae, respectively. (**h**–**i**) Volcano plots depicting differentially expressed genes at 8.5 and 12 hpf in DMH1 vs control embryos treated from 4 to 8 hpf. anim animal, dor dorsal, lat lateral. Drawings are not to scale.

We therefore analysed the expression of core BMP pathway components during cleavage and larval stages. Our expression profile analysis reveals that the genes *bmp2/4*, *smad6*, and *dlx* (an alternative readout of BMP signalling^45^ given the lack of crossreactivity for the pSMAD1/5/8 antibody in *P. dumerilii*) start to be expressed from 6 to 8 hpf in dorsal cell lineages of the head and trunk, including 2d and its progeny, as well as the progeny of 1c and 1d (Fig. 4b; Extended Data Fig. 4a, c). On the opposite side, the BMP antagonists *noggin* and *bmp3* are expressed ventrally in both the head and trunk (Fig. 4c; Extended Data Fig. 4b, d). Their expression patterns correspond to those of known ventral head (e.g., *nk2.1*, *hnf6*) and trunk markers (e.g., *gsc*, *foxA*, and *bra*) (Extended Data Fig. 4e–g). Therefore, the expression of *bmp2/4* and putative BMP target genes on the dorsal side, in contrast to the ventral expression of genes encoding BMP antagonists, strongly suggests that the BMP pathway could play a role in DV axis specification during the early unequal cell divisions in *P. dumerilii*.

To investigate this hypothesis, we inhibited and overactivated the BMP signalling with a continuous treatment of DMH1 and rBMP4, respectively, from 2 hpf. DMH1 treatment induced the ventralisation of *P. dumerilii* larvae at 72 hpf, which lacked both larval and adult eyes, the dorsoanterior ciliary band, and the nephroblasts (Fig. 4d). These ventralised larvae exhibited radialisation of gland cells expressing *hnf6,* a gene usually restricted to the ventral region in wild-type larvae (Fig. 4e). To refine the temporal window of BMP signalling activity essential for DV patterning, we conducted staggered DMH1 treatments between 2 and 12 hpf. This revealed a critical period of BMP signalling activity around 6 hpf, when the embryo consists of ∼16 cells (Extended Data Fig. 5a). Indeed, inhibition of the BMP signalling from 4 to 8 hpf (8 to ∼66-cell stages) leads to larvae with ventralised heads, as in continuous DMH1 treatment (Fig. 4d, e; Extended Data Fig. 5b,c). The ventralisation induced by the BMP signalling inhibition can already be observed at 12 hpf with the loss of dorsal markers, including *bmp2/4*, *smad6,* and *dlx* and the ectopic expression of ventral head genes such as *noggin*, *bmp3*, and *nk2.1* throughout the entire head region (Fig. 4f, g). Nephroblasts and the akrotroch (a dorsal circlet of cilia in the head), originally formed from a pair of large cells in the dorsal head^43,46,47^, also disappear, as revealed by the absence of expression of *fzCRD1* and *tektin2*, respectively (Extended Data Fig. 5d, e). In contrast with the DMH1 treatment, rBMP4 exposure induced the dorsalisation of larvae at 72 hpf, which lost the ventral gland cells in 24 and 72 hpf (Fig. 4d, e), and did not express ventral markers (*noggin* and *nk2.1*). In addition, overactivation of the BMP signalling led to the ectopic expression of dorsal genes (*bmp2/4*, *dlx*, and *smad6*) into the ventral head territory at 12 hpf (Fig. 4f, g). Unexpectedly, both DMH1 and rBMP4-treated larvae show only mild phenotypes in the trunk region. BMP signalling inhibition results in a slightly bent trunk at 72 hpf (Fig. 4d; Extended Data Fig. 5b), reminiscent of the phenotype observed in *C. teleta* after BMP inhibition^16,48^. The trunk expression of dorsal markers in the 2d (*bmp2/4*, *dlx*, *smad6*) and 4d progeny (*brachyury*) was also reduced (Fig. 4f; Extended Data Fig. 5f), while the ventral trunk markers *foxA* and *goosecoid* show a dorsal expansion in DMH1-treated 12 hpf embryos (Extended Data Fig. 5f). Ectopic BMP activation affected the patterning of the trunk even less, except for *goosecoid* expression and the alteration of the stomodaeum anlage (Fig. 4d; Extended Data Fig. 5g). Altogether, our findings demonstrate that BMP signalling is essential for DV patterning during early spiral cleavage in *P. dumerilii*. However, its effect is primarily restricted to specifying dorsal cell lineages in the head region.

To capture the early and late transcriptional responses induced by BMP signalling during head DV patterning, we performed comparative bulk transcriptomics analysis of control and DMH1-treated embryos (from 4 to 8 hpf) at four developmental time points (8.5, 12, 18, and 24 hpf) (Extended Data Fig. 5h, i; Supplementary Tables 18–26). At 8.5 and 12 hpf, genes orthologous to the vertebrate BMP synexpression group^49^, including *vent, bambi, smad6*, *dlx, tbx3,* and *id4*, as well as transcription factors like *hes-related* factors, *hmx3*, and *bsh*, and similar to *O. fusiformis*, the ligand *delta* were amongst the significantly downregulated genes in DMH1-treated embryos (Fig. 4h; Supplementary Tables 19, 21). In contrast, ventral genes such as *nk2.1*, *noggin* and *bmp3*, which expand their expression after DMH1 treatment (Fig. 4g), are significantly upregulated at these time points (Supplementary Tables 20, 22), supporting the notion that in the absence of BMP signalling, the head ventralisation is the default state for the ‘dorsal’ micromeres. Notably, genes like *smad6, dlx, bambi, tbx3,* and *vent* are downregulated in both embryonic and larval stages, while *nk2.1* is upregulated (Fig. 4h, i; Extended Data Fig. 5j, k; Supplementary Tables 19-26). Remarkably, only *delta* and *dlx* are shared downregulated genes between the early DMH1-treated embryos of *O. fusiformis* (6 hpf) and *P. dumerilii* (8.5 hpf). In the later larval stages, genes such as *bsh, gcm, six1*, *vsx2*, and *gata1/2/*3 were amongst the significantly downregulated genes, whereas *hnf6* and *ptf1* among the significantly upregulated genes (Extended Data Fig. 5j, k; Supplementary Tables 23-26), supporting their roles in the development of the dorsal and ventral head features, like the dorsoanterior brain region and the ventral gland cells, respectively. Therefore, our results demonstrate that a short early BMP signalling event is sufficient to establish a transcriptional DV asymmetry in the head of the *P. dumerilii* embryo. However, the transcriptional response after BMP signalling in *P. dumerilii* differs considerably from that observed in *O. fusiformis*, at least in its initial stages.

### The 2d blastomere is the head DV organiser in P. dumerilii

In the autonomously developing annelid *C. teleta*, the 2d cell functions as a head axial organiser^15,16,50^. Given the strong expression of *bmp2/4* in the 2d cell (Fig. 4b) during the critical timing of BMP signalling, we hypothesised that a BMP-mediated signal from this cell could induce the BMP-mediated DV patterning of the head. To test this, we ablated the 2d cell, as well as its progenitor, the 1D blastomere, its sister cell, the 2D blastomere, and its early progeny (Fig. 5a; Extended Data Fig. 6a). As expected, only the ablation of the 2d cell and its progenitor phenocopied the DV patterning and differentiation defects observed in the head observed in DMH1-treated embryos (Fig. 4d, e; Fig. 5b, c; Extended Data Fig. 6b, c).

**Figure 5.**
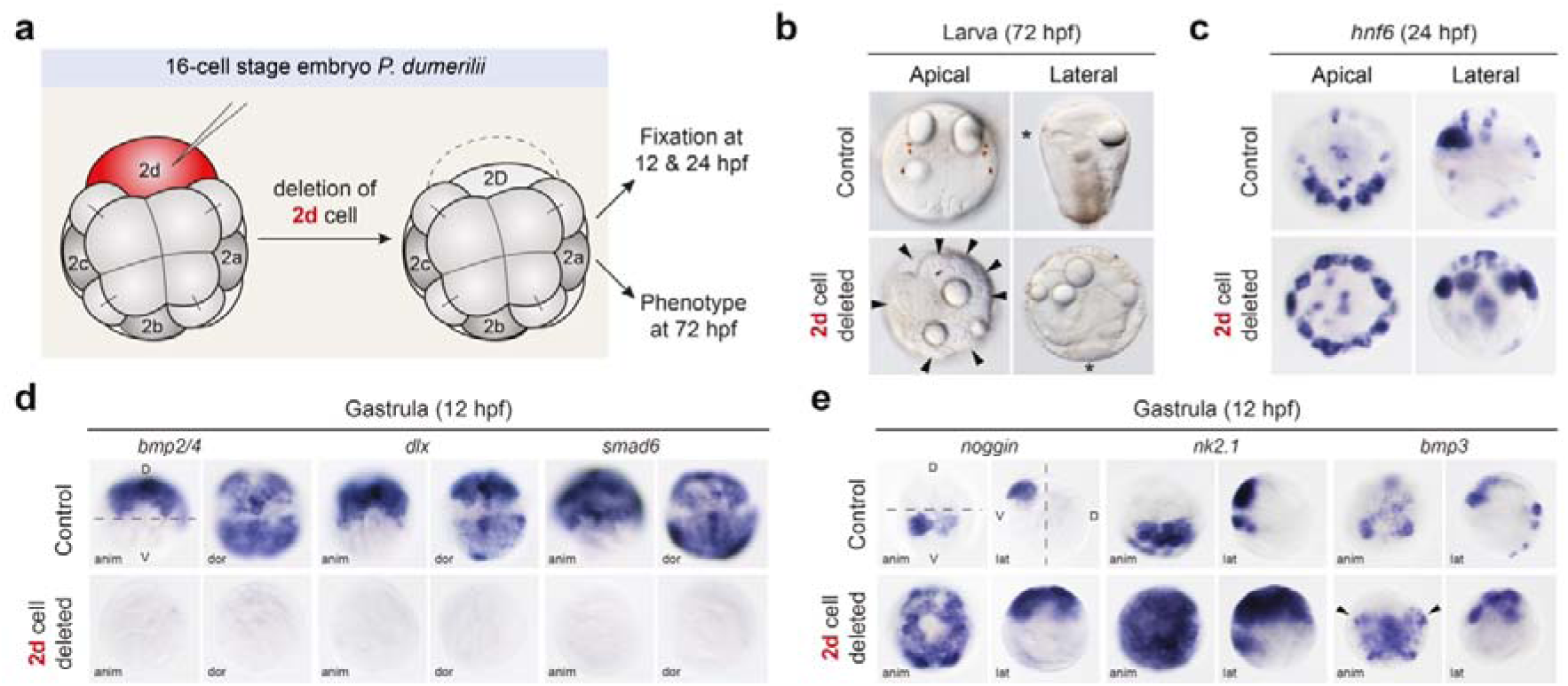
2d functions as the head embryonic organiser in *P. dumerilii*. (**a**) Diagram of the 2d-cell (red) ablation in the 16 cell-stage embryo, and subsequent experimental procedures. (**b**) Differential Interference Contrast (DIC) images of 72 hpf larvae after 2d ablation. Note the absence of bilateral eyes, the radialisation of the larval glands (arrowheads) and the presence of the mouth (asterisk) more posterior in comparison with controls. (**c**–**e**) Whole mount *in situ* hybridisation of dorsoventral markers in controls and embryos from 2d- cell ablations at (**c**) 24 hpf and (**d**–**e**) 12 hpf embryos. Ablation of the 2d cell phenocopies DMH1 treatments (compare to Fig. 4f–h). anim animal, dor dorsal, lat lateral.

Importantly, exposure to rBMP4 protein partially rescued this ventralised phenotype (Extended Data Fig. 6d). In 2d-deleted embryos, the expression of the dorsal markers *bmp2/4*, *dlx*, and *smad6* was abolished, whereas the ventral markers *noggin*, *bmp3*, and *nk2.1* expanded dorsally in the head (Fig. 5d, e). The ablation of the 2d cell also abolished the expression of *bmp2/4*, *dlx*, and *smad6* in the trunk (Fig. 5d), suggesting that the dorsal trunk domain expressing these genes derives directly from the 2d cell. However, while the expression of *brachyury* and *foxA* is mostly retained in the 2d deleted embryo at 12 hpf, *foxA* expression and mouth position are strongly affected in the 2d deleted 72 hpf larvae (Extended Data Fig. 6e, f). Therefore, in the early embryo of *P. dumerilli*, the 2d cell, expressing *bmp2/4* from 6 hpf, acts as an embryonic organiser responsible for the DV patterning of the head.

This organiser, via BMP signalling, activates the expression of dorsal head genes, including *bmp2/4* itself, and restricts ventral head genes such as *noggin* or *bmp3* in the ventral head region. After differentiation, these two regions along the DV axis will give rise to gland cells on the ventral side, and adult and larval eyes on the dorsal side of the larval head (Extended Data Fig. 6g).

### Activin/Nodal affects dorsoposterior development in O. fusiformis

The transition to a DV specification relying on Activin/Nodal signalling in some annelids with autonomous spiral cleavage raises the question of the ancestral role of this pathway in annelids. To address this, we first explored the Activin/Nodal pathway in *O. fusiformis*. This annelid has three Activin/Nodal/inhibin ligands —*inhibin-like 4*, *inhibin-like 5,* and *Nodal*–– and two activin receptors — *ACVR1* and *ACVR2*^25^. While *inhibin* ligands are mainly expressed in late development, *Nodal* expression peaks at 5 hpf, when it is strongly detected in the gastral plate, including the embryonic organiser (Extended Data Fig. 7a–b). With gastrulation, *Nodal* becomes expressed solely in the embryonic organiser (Extended Data Fig. 7b). At these stages the *ACVR2* receptor and the secondary messenger *smad2/3* are weakly but ubiquitously expressed (Extended Data Fig. 7b). To uncover the role of Activin/Nodal signalling, we treated embryos of *O. fusiformis* with the specific Nodal inhibitor SB431542 and recombinant Activin-A at the time of axial patterning (Extended Data Fig. 7c–d; Supplementary Table 3–4). Treatment with SB431542 resulted in smaller larvae (70%) with a smaller chaetal sac and shorter chaetae (Extended Data Fig. 7e, f). Notably, SB431542 prevents neither the formation of the organiser nor the activation of pSMAD1/5/8, and the treated larvae retain a normal expression of oral and hindgut markers (Extended Data Fig. 7g, h; Supplementary Table 5). However, after rActivin A treatment, larvae phenocopy the rBMP4-treated larvae, exhibiting ectopic chaetal sacs and loss of *gsc* expression (Extended Data Fig. 7e, h; Supplementary Table 3–5). Therefore, Activin/Nodal is unnecessary to establish the DV polarity in *O. fusiformis*. Yet, the inhibition of this pathway is reminiscent of the effects of inhibiting the BMP signalling.

To discern if the effect of Activin/Nodal on DV polarity relies on similar genes downstream of BMP signalling, we performed comparative transcriptomic profiling of control and SB431542 and rActivin A-treated embryos (Extended Data Fig. 8a, b). We found 27 upregulated and 41 downregulated genes in SB431542-treated embryos compared to the controls (Extended Data Fig. 8c, d; Supplementary Table 7, 27–28). Surprisingly, SB431542 and DMH1-treated embryos do not share any downstream genes despite leading to comparable (but weaker in SB431542) phenotypes in the larvae (Extended Data Fig. 8e; Supplementary Tables 7–11). Amongst the downstream genes, we focused on the three transcription factors with a known developmental role: *otx*, expressed in precursors of the ciliary band and later the gut^23^; *hb*, expressed in the gastral plate and endoderm; and *prospero*, expressed in the ectoderm and oesophagus (Extended Data Fig. 8d, f,g). –h). In all cases, their expression is dramatically downregulated in the blastula after SB431542 treatment (Extended Data Fig. 8h). Unlike SB431542, treatment with rActivin A leads to many differentially expressed genes, including genes expressed in the apical organ (e.g., *six3/6* and *rx1*) and internalised cells (e.g., *zag1*) (Extended Data Fig. 9a, b; Supplementary Table 12). However, only a small fraction of differentially expressed genes (∼3%) were shared with those affected by rBMP4 treatment (Extended Data Fig. 9c; Supplementary Tables 29–30). This included five downregulated (*osr*, *foxG*, *foxA*, *fojo*, and *cbc1*) and two upregulated (*wnt5* and *msx*) genes (Extended Data Fig. 9e–f; Supplementary Table 12).

Altogether, our findings show that the BMP and Activin/Nodal pathways largely regulate distinct sets of genes, suggesting that the effect of Activin/Nodal in dorsoventral development is likely secondary and/or indirect in *O. fusiformis*.

### The Activin/Nodal pathway does not regulate DV patterning in P. dumerilii

In annelids with autonomous development, such as *C. teleta* and *C. pergamentaceus*^15,16,19^, a reduced role of the BMP signalling in DV patterning co-occurs with a compensatory and more prominent axial patterning function of the Activin/Nodal pathway. To determine if this shift also occurs in *P. dumerilii*, we treated embryos continuously with the inhibitor SB431542 at concentrations of 10 and 20 µM from 2 hpf (Extended Data Fig. 10a, b). At 72 hpf, treated larvae displayed normal morphology, with a correct DV patterning of both head and trunk (Extended Data Fig. 10a). However, we note the absence of adult eyes at 20 µM in the dorsal head region (Extended Data Fig. 10a). Accordingly, the expression of the ventral head marker *nk2.1*, which expands dorsally after BMP inhibition (Fig. 4g), shows a normal expression in SB431542-treated embryos (Extended Data Fig. 10b), further supporting that the DV patterning of the head is not affected after inhibiting the Activin/Nodal signalling pathway, unlike in *C. teleta*^15,16^. Interestingly, Nodal shows a left-right asymmetric expression in the 24 hpf larva of *P. dumerilii*, when it is expressed on the right side and partially depends on normal BMP signalling (Extended Data Fig. 10c). Therefore, unlike *O. fusiformis* and *C. teleta*, the Activin/Nodal pathway is not required for DV patterning in *P. dumerilii*.

### DV development relies on different downstream genes in O. fusiformis and C. teleta

An evolutionary transition to Activin/Nodal-mediated DV patterning might have rewired this signalling pathway to control axial polarity genes originally downstream of BMP signalling. To assess this hypothesis, we first compared the temporal expression profile of the thirteen one-to-one orthologous developmental genes between *O. fusiformis* and *C. teleta*^18^ that are downregulated by DMH1 and SB431542 and upregulated by rBMP4 in *O. fusiformis*. None of them show a pattern indicating upregulation after the organising activity of Nodal at the 16-cell stage in *C. teleta*^50^ (Extended Data Fig. 10d), except for *otx* and *fojo*, which are broadly expressed in the animal pole at that time^51,52^. Considering that transcriptomic dynamics might not fully reflect the spatiotemporal expression of these genes, we investigated the expression in *C. teleta* of three genes under different regulatory conditions in *O. fusiformis*: *fer3*, activated by BMP; *osr*, inhibited by BMP; and *hb*, activated by Nodal.

Like in *O. fusiformis*, *fer3* is expressed in the chaetoblasts, *osr* marks the boundary between the foregut and the midgut, and *hb* is expressed in the developing brain and ventral nerve cord in *C. teleta* (Extended Data Fig. 10e). However, inhibition of DV polarity with SB431542^16^, which also affects trunk elongation in *C. teleta* (Supplementary Table 31), restricts, but does not eliminate, the expression of *osr* and *hb* to the anterior body (Extended Data Fig. 10f, g), suggesting that these are not direct downstream genes of Activin/Nodal in this annelid. Therefore, our data indicate that changes in the upstream signalling that specifies the DV axis rewired the downstream effector genes between *O. fusiformis* and *C. teleta*.

## Discussion

Combining a multispecies approach with comparative transcriptomics, blastomere deletions, and pharmacological inhibition of the ERK, BMP and Activin/Nodal signalling pathways, our study clearly reveals a conserved use of the ERK1/2 and BMP pathways to specify the DV axis in distantly related annelids with conditional spiral cleavage (Fig. 6a). This is similar to other spiralians like molluscs^9,12,24^, particularly the gastropod with conditional spiral cleavage *Lottia peitaihoensis*, which also uses FGFR, ERK1/2 and BMP to define the embryonic organiser and bilateral symmetry^12,24^. In contrast, the anti-neural role of the BMP pathway, as seen in vertebrates^3^ and arthropods^4^, remains unclear across spiralians. In some molluscs, BMP promotes neurogenesis^9,12^, while in others (e.g., *Crepidula fornicata*^13^) and in annelids (e.g., *C. teleta*, *Helobdella* and *P. dumerilii*), this signalling pathway acts locally to set up the DV coordinates in the neuroectoderm^14,17,48,53^. In *O. fusiformis*, BMP signalling disrupts the patterning of the nervous system, but mature neurons (as seen through neuropeptides such as RYamide) are still present. Therefore, an early role of the BMP signalling pathway in setting up the DV axis is ancestral to Spiralia, as in most other bilaterians^5^ (Fig. 1a, 6a). Yet, unlike in other non-spiralian lineages, this axial function uniquely occurs downstream of the FGFR- ERK1/2-mediated patterning system and is likely decoupled from a potential neurogenic role later in development in Spiralia.

**Figure 6.**
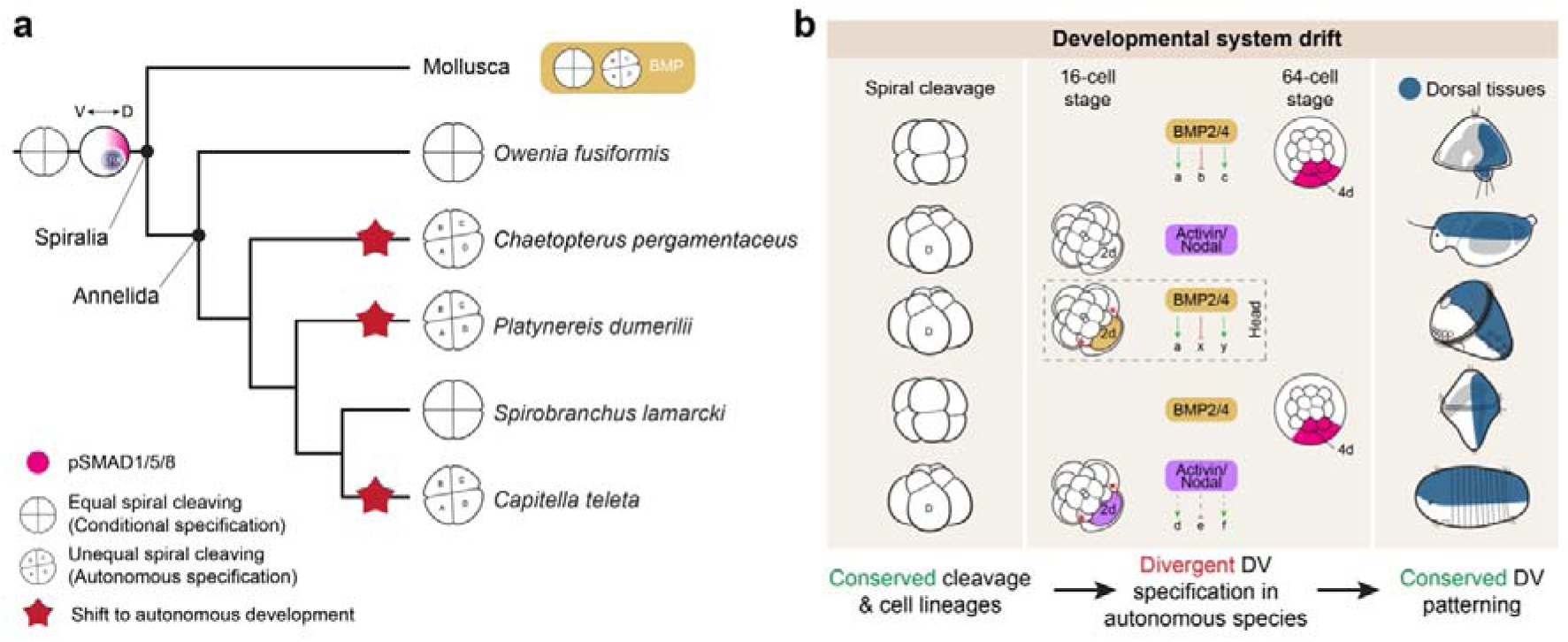
The ancestral role of BMP in Spiralia and the evolution of developmental system drift in dorsoventral patterning. (**a**) Evolutionary scenario for the developmental role of the BMP pathway in axial patterning in Mollusca and Annelida and its last common ancestor (assumed to be the last common ancestor to Lophotrochozoa based on current phylogenies). Given BMP’s role in DV patterning and the presence of a dorsoventral gradient of pSMAD1/5/8 in molluscs and annelids with conditional/equal spiral cleavage, a ERK1/2- BMP signalling axis defining the posterodorsal side of the embryo is likely ancestral to Spiralia, absent in species that have transitioned to an autonomous/unequal mode of spiral cleavage (red stars). (**b**) Schematic drawings representing the cellular and molecular specification of the dorsoventral axis in the studied annelid species. All these annelids exhibit a conserved and homologous cleavage programme, cell lineages and dorsoventral axes. However, the cells and molecular underpinnings can diverge profoundly, a phenomenon termed developmental system drift, in species that have transitioned to autonomous development. While annelids with conditional spiral cleavage employ ERK1/2 upstream of BMP to specify the bilateral symmetry and dorsoventral axis, respectively, species with autonomous specification define the axial polarity and dorsoventral axis around the 16-cell stage, with a prominent role for the 2d blastomere, at least in *P. dumerilii* and *C. teleta*. Moreover, divergent upstream regulators trigger dorsoventral specification, with Activin/Nodal as the primary regulator in *C. pergamentaceus* and *C. teleta*, and the BMP pathway driving dorsoventral polarity only in the head in *P. dumerilii*. The divergence in upstream regulators is also concomitant with divergence in downstream effector genes. Drawings are not to scale.

An ancestral FGFR-ERK1/2-BMP dorsoventral patterning implies that distantly related annelids with autonomous specification have independently diverged to use Activin/Nodal for DV specification or restricted BMP signalling to DV patterning in only a subset of tissues, such as the head^53^ or the local mid-line ectoderm^17,53^. Importantly, in both *C. teleta*^15,16,50^ and *P. dumerilii*, the same blastomere, the 2d cell, exerts this early axial organising role, albeit through Activin/Nodal and BMP signalling, respectively (Fig. 6b). Given the conservation of cell lineages and fate maps across annelids and spiralians, a homologous body region (i.e., the DV tissue) is thus specified through different genetic and developmental mechanisms in annelids, a phenomenon called developmental system drift (DSD)^54,55^. DSD is widespread yet poorly understood in animals and reflects the variability of embryogenesis, gene regulatory networks, and morphogenic events^54^. Not surprisingly, DSD was previously invoked regarding DV specification among spiralians^11,15^, given the dramatically different role of BMP in DV specification between molluscs and autonomous annelids. Our study refines this scenario, indicating that DSD in DV patterning co-occurs with transitions to an autonomous mode of spiral cleavage in annelids (Fig. 6b) and possibly in some molluscan lineages (e.g., *Crepidula fornicata*).

How could DSD occur, and what are its implications at the gene regulatory level? In *O. fusiformis*, Activin/Nodal influences DV axis development, a condition reminiscent of deuterostomes^56–58^, in an ERK1/2-independent manner (Fig. 6b). A transition to an autonomous specification of the axial identities through the asymmetric inheritance of maternal determinants^59^ bypasses the axial patterning role of ERK1/2, thereby eliminating the upstream regulator of the BMP signalling to specify the DV axis. This might favour the Activin/Nodal pathway gaining a more predominant role in DV patterning in some annelids, while repurposing through yet unknown maternal cues, the BMP signalling predominantly for the anterior/head neuroectodermal regionalisation^53^ in others. Notably, our transcriptomic data in *O. fusiformis* and *P. dumerilii* and comparison with *C. teleta* suggest that divergence in DV patterning upstream regulators likely involves significantly different effector genes, even when the cells contributing to each body region are conserved. This indicates that DSD does not rewire upstream signalling to activate a conserved downstream cascade but likely implies recruiting different networks for the specification of the DV axis (Fig. 6b).

Altogether, our work solves current questions on the ancestral role of the BMP signalling pathway in DV axis specification in Annelida and Spiralia, providing a conceptual framework and tractable systems to investigate the evolutionary and developmental mechanisms generating DSD during animal embryogenesis.

## Methods

### Animal husbandry and specimen collections

Adults, embryos, and larvae of *O. fusiformis*, *P. dumerilii*, *C. teleta* and *S. lamarcki* were collected as described before^32,39,46,60,61^. Embryos of *O. fusiformis* and *P. dumerilii* used for RNA sequencing were snap-frozen in liquid nitrogen and stored at –80 °C. Embryos and larvae of *O. fusiformis*, *S. lamarcki* and *C. teleta* used for other analyses were fixed in 4% paraformaldehyde in artificial sea water (ASW), and stored in either 1x phosphate-buffered saline (PBS) with sodium azide at 4°C or methanol at –20 °C. Embryos and larvae of *P. dumerilii* for whole-mount *in situ* hybridisation were fixed in 4% paraformaldehyde in PBS for one hour at room temperature and stored in methanol at –20°C.

### Drug treatments

U0126 (Promega; Cat No.: V1121 or Merck-Sigma; Cat No.: 19-147), DMH1 (Merck-Sigma; Cat No.: D8946) and SB 431542 hydrate (Merck-Sigma; Cat No.: S4317) 10 mM drug stocks were made in dimethyl sulfoxide (DMSO). Stocks of 100 μg/ml recombinant zebrafish BMP4 protein for *O. fusiformis* (R&D; Cat No.: 1128-BM), recombinant human BMP4/BMP7 heterodimer for *P. dumerilii* (R&D; Cat No.: 3727-BP), and recombinant human/mouse/rat Activin A protein (R&D; Cat No.: 338-AC) were made in 4mM HCl + 0.1% BSA and stored at –80°C. Working solutions of the treatments were prepared in ASW to the desired concentrations, using equivalent volumes of solvents (DMSO or 4mM HCl + 0.1% BSA) as negative controls.

### RNA-seq profiling and differential gene expression analyses

For *O. fusiformis*, total RNA was extracted from 6 hpf blastulae right after treatments using the Monarch Total RNA Miniprep kit (New England Biolabs; Cat No.: #T2010) and used to prep strand-specific mRNA Illumina libraries that were sequenced at the Oxford Genomics Centre (University of Oxford, UK) over one lane of an Illumina NovaSeq6000 system in 2 × 150 bases mode. Adapter and low-quality bases were removed using FastP v.0.20.1^62^.

Cleaned reads were mapped to the *O. fusiformis* genome annotation^18^ using Kallisto v.0.46.2^63^. Read counts normalisation and differential gene expression analyses for each pairwise comparison between control and treatment conditions were performed with the R package DESeq2 v.1.26.0 ^64^. The enrichment of Gene Ontology terms in the differentially expressed genes was calculated with the TopGO R package^65^.

For *P. dumerilii*, RNA samples were collected at 8.5, 12, 18, and 24 hpf. A total of six biological batches were collected, each comprising eight samples representing the four developmental stages for DMSO- and DMH1-treated embryos and larvae. The top three batches were selected for RNA sequencing based on (i) developmental success rate, (ii) signal clarity and reproducibility in gene expression analyses, and (iii) RNA quality. RNA extraction was performed using TRIzol® reagent (Invitrogen; Cat No.: 15596026), following the manufacturer’s instructions. Strand-specific mRNA Illumina libraries were prepped and sequenced at the NGS core facility in Academia Sinica (Taiwan) over one lane of an Illumina^®^ HiSeq 2500 System. Raw reads were quality-trimmed to remove adapters and low-quality bases using Trimmomatic v.0.39^66^. Trimmed reads were mapped to the *Platynereis* reference genome v.0.2.1 using STAR v.2.7.11b^67^, and gene abundance was estimated with featureCounts v.2.0.1^68^. Read counts normalisation and differential gene expression analyses for each pairwise comparison between control and treatment conditions were performed with the R package DESeq2 v.1.26.0^64^, adjusting log2 fold changes with the shrinkage approach included in DESeq2.

### Gene isolation and gene expression analyses

For *O. fusiformis* and *C. teleta*, candidate genes and phenotypical marker genes were amplified as previously described^7^ using gene-specific primers and a T7 adaptor. Riboprobes were synthesised with the T7 MEGAscript kit (ThermoFisher; Cat No.: AM1334) and stored at a concentration of 50 ng/μl in hybridisation buffer at –20 °C. Whole-mount colourimetric and fluorescent *in situ* hybridisation in embryonic and larval stages were conducted as described elsewhere for *O. fusiformis* ^32^ and *C. teleta* ^60^. For *P. dumerilii*, DIG-labelled antisense RNA probes were synthesised using the DIG RNA Labelling Mix (Roche, Cat. No: 11277073910) and whole mount colourimetric in situ hybridisation was performed as described elsewhere^46^.

### Whole-mount immunohistochemistry

Whole-mount immunohistochemistry, nuclear and F-actin staining, were conducted as described previously^7,31,32^. The primary antibodies used were mouse anti-acetylated α-tubulin (clone 6-11B-1, Merck-Sigma; Cat No.: #MABT868, 1:500), mouse anti-β-tubulin (E7, Developmental Studies Hybridoma Bank), and *P. dumerilii-*derived rabbit anti- RYamide^31,33,34^ diluted in 5% normal goat serum (NGS) in 1x PBS + 0.1% Triton X-100 (PTx) overnight. After several PTx washes, animals were incubated in the following secondary antibodies: 1:800 goat anti-mouse AlexaFluor 488 (ThermoFisher Scientific; Cat No.: A32731), 1:800 goat anti-mouse AlexaFluor 647 (ThermoFisher Scientific; Cat No.: A- 21235), and 1:800 goat anti-rabbit AlexaFluor 555 (ThermoFisher Scientific; Cat No.: A32731). To detect the activation of SMAD1/5/8, the primary antibody rabbit anti-Phospho- Smad1/5 (Cell Signaling Technology; Cat No.: 9516) was used following a modified protocol used in other spiralians^23^. Briefly, wild type, control and treated embryos stored in methanol were rehydrated gradually with 1x PBS + 0.1 % Tween-20, and permeabilised for one hour with 1x PBS + 0.2% Tween-20/0.2% Triton X-100 (PTwTx). Embryos were then blocked for one hour with 5% NGS in PTwTx and incubated overnight in 1:10 anti-pSmad1/5. After several washes with PTwTx, the embryos were incubated with 1:800 anti-rabbit AlexaFluor 555 (ThermoFisher Scientific, cat#: A-21428) plus 4′,6-diamidino-2-phenylindole (DAPI, ThermoFisher Scientific, #D3571, stock 2mg/ml, 1:2000) as a nuclear marker overnight and stored in 70% glycerol in PBS. The same primary antibody against Phospho-Smad1/5 was used in *P. dumerilii*, producing no reliable staining. For dp-ERK1/2 staining in *S. lamarcki*, we use embryos stored in methanol at –20 °C and a 1:100 dilution of the primary antibody mouse anti-dp-ERK1/2 (Merck-Sigma; Cat No.: #M9692) as described before^7^.

### Phenotype characterisation and classification

For *O. fusiformis*, we used morphological characters, such as the presence/absence of chaetal sacs, and gene expression to determine the loss or gain of dorsoposterior and neural tissue in the larva. For *S. lamarcki,* we used the presence/absence of the hyposphere (future trunk of the worm) to determine the phenotype of the drug inhibitions. With *P. dumerilii*, morphological phenotypes were scored in 72 hpf larvae, which were anaesthetised using fresh fixative buffer and placed in a petri dish (35 mm diameter) for motion-capture imaging using an Axiocam 506 colour camera in conjunction with a Zeiss Axioskop microscope.

Morphological scoring included adult eye number, body shape, and overall development.

### Imaging

Samples were mounted in 70% glycerol in 1x PBS. DIC images of the colourimetric *in situs* were obtained with a Leica 560 DMRA2 upright microscope equipped with an Infinity5 camera (Lumenera) or a Zeiss Axioskop microscope. Immunohistochemistry images were acquired with a Leica SP5 Laser Scanning Confocal, a Leica Stellaris 8, or a Nikon CSU-W1 Spinning Disk Confocal. Z projections were analysed with Fiji, and brightness/contrast was adjusted with Adobe Photoshop (v22.2.0). Figs were designed with Adobe Illustrator (v28.6).

## Acknowledgements

We thank members of the Martín-Durán, Schneider and Ferrier Labs, Elaine Seaver, and technical staff within the Department of Biology at Queen Mary University of London for their feedback and support. This study was funded by the European Research Council (Starting Grant, Action number 801669) to JMMD, the Biotechnology and Biological Sciences Research Council LIDo Summer Research Placement to AMCB, the Biotechnology and Biological Sciences Research Council (BB/Y004221/1 and BB/Y004868/1) to AMCB, DEKF and JMMD, and by the National Science Foundation, USA (NSF IOS-1455185), National Science and Technology Council, R.O.C. (MOST 108-2311-B-001-002-MY3), and Academia Sinica (AS-CDA-110-L02 & intramural funds) to SQS. We thank the Oxford Genomics Centre at the Wellcome Centre for Human Genetics (funded by Wellcome Trust grant reference 203141/A/16/Z) for generating and initially processing the sequencing data. This research utilised Queen Mary’s Apocrita HPC facility, supported by QMUL Research-IT, and the confocal microscope supported by BBSRC grant BB/W019698/1.

## Authors contributions

AMCB, EH, SQS and JMMD conceived and designed the research. AMCB, EH, SMM, YL, DEKF and JMMD performed collections. AMCB, EH and YL performed the extractions.

AMCB, YL, TL and JMMD performed all RNA-seq analyses. AMCB, EH, and SMM performed gene expression analyses, with help from IL and AP. AMCB, SQS, and JMMD drafted the manuscript; all authors critically read and commented on it.

## Data availability

All expression data generated in this manuscript has been uploaded to SRA (project ID PRJNA1267570) and Gene Expression Omnibus (study ID GSE297936).

---

**Extended Data Fig. 1.**
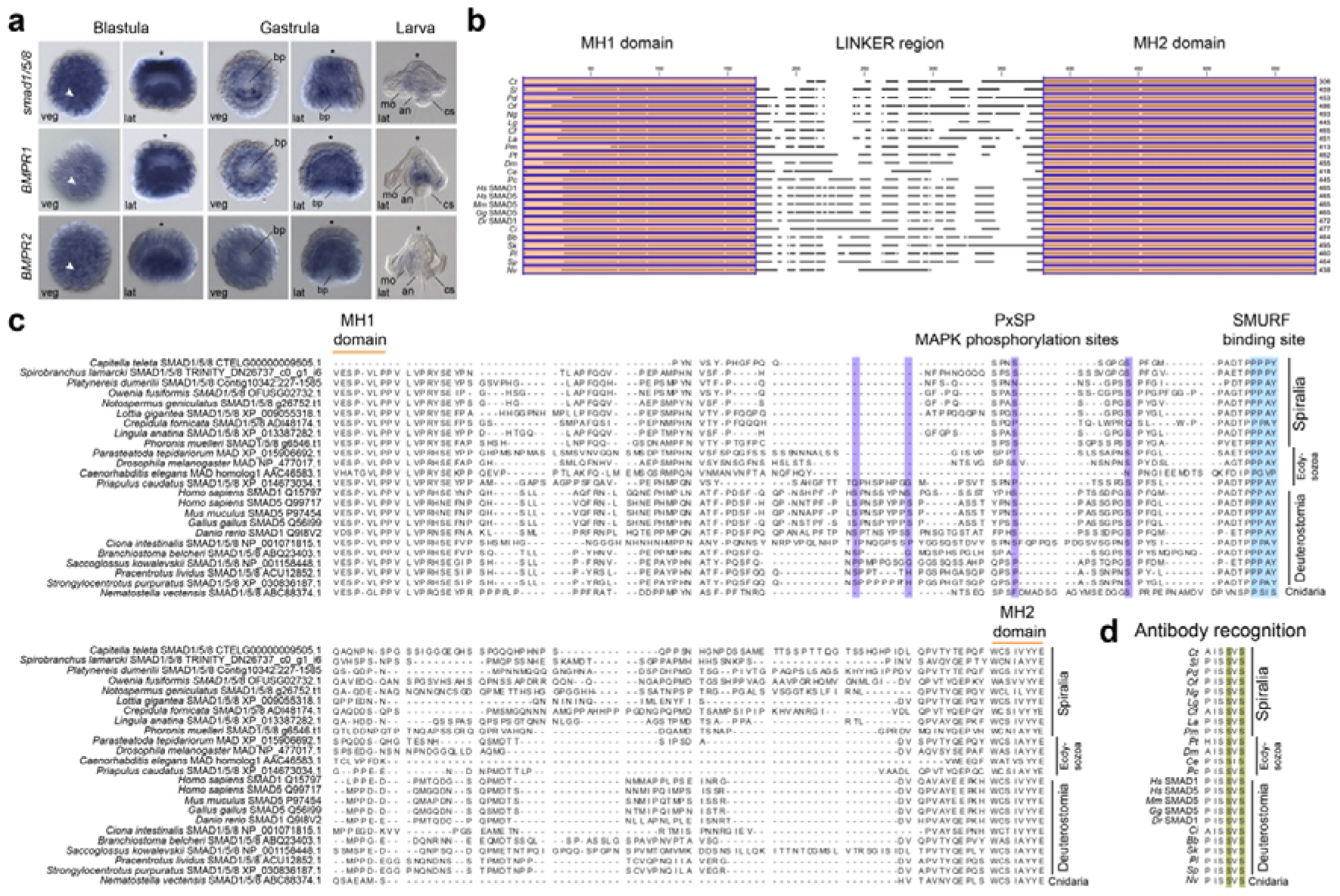
The BMP pathway and conservation of Smad1/5/8 in *O. fusiformis*. (**a**) Whole mount *in situ* hybridisation of the secondary messenger *smad1/5/8* and receptors *bmpr1* and *bmpr2* of the BMP pathway at the blastula (6 hpf), gastrula (9 hpf) and larval (24 hpf) stages in *O. fusiformis*. (**b**) Schematic diagram of a multiple protein alignment of the MH domains and linker region of SMAD1/5/8 across bilaterian and cnidarians highlighting the high conservation of the MH domains. (**c**) Multiple protein alignment in the linker region of SMAD1/5/8 showing the presumptive MAPK phosphorylation sites (highlighted in violet). Only Deuterostomes have all sites conserved. (**d**) Multiple protein alignment of the C-terminus of SMAD1/5/8 proteins showing the conserved residues recognised by the pSMAD1/5/8 antibody used in this study. In (**a**), the arrowheads point to the 4d organiser, and the asterisks mark the animal/apical pole. Scale bars are 50 µm. an anus, ao apical organ, bp blastopore, ch chaetae, cs chaetal sac, fg foregut, lat lateral, mo mouth, pt prototroch, veg vegetal.

**Extended Data Fig. 2.**
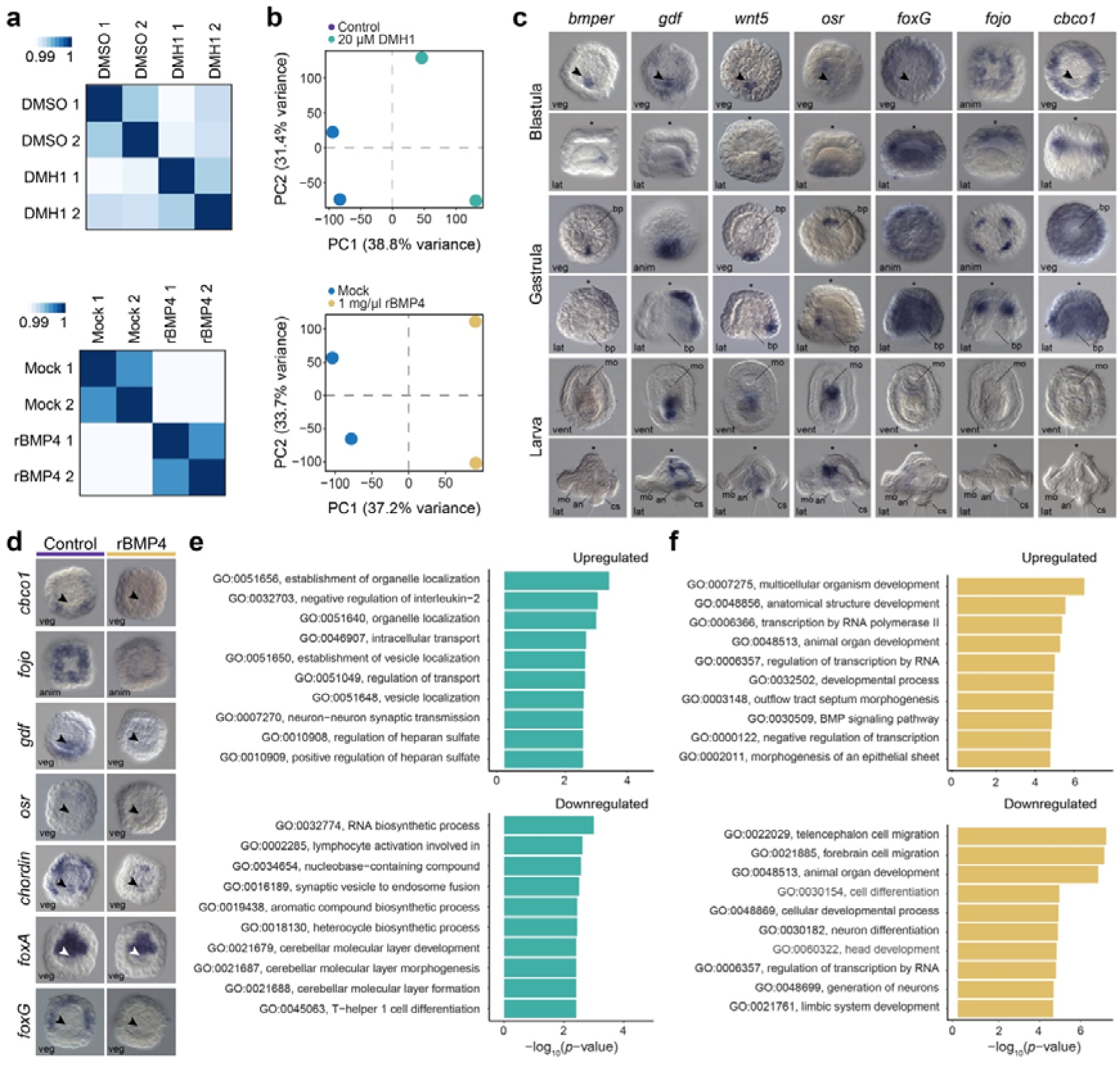
Differential gene expression analyses in DMH1 and rBMP4- treated *O. fusiformis* embryos. (**a**) Hierarchically clustered pairwise correlation matrix between DMH1 (top) and rBMP4 (bottom) treated and control samples. The scale shows the relative difference in Euclidean distance. (**b**) Principal component (PC) analysis plots for DMH1 (top) and rBMP4 (bottom) transcriptomic analyses indicate that the main source of transcriptional variability between samples is the treatment condition (PC1). (**c**) Whole mount *in situ* hybridisation of selected candidate, differentially expressed genes in the rBMP4- treated condition, at the blastula, gastrula, and larval stages. (**d**) Whole mount *in situ* hybridisation validates the downregulation of candidate genes at the blastula stage (6 hpf) after rBMP4 treatment. (**e**, **f**) Bar plots showing the top ten enriched Gene Ontology terms in differentially expressed genes after DMH1 (**e**) and rBMP4 treatment (**f**). In (**c**, **d**), the arrowheads point to the 4d organiser, and the asterisks mark the animal/apical pole. Scale bars are 50 µm. an anus, anim animal, ao apical organ, bp blastopore, ch chaetae, cs chaetal sac, fg foregut, lat lateral, mo mouth, pt prototroch, veg vegetal.

**Extended Data Fig. 3.**
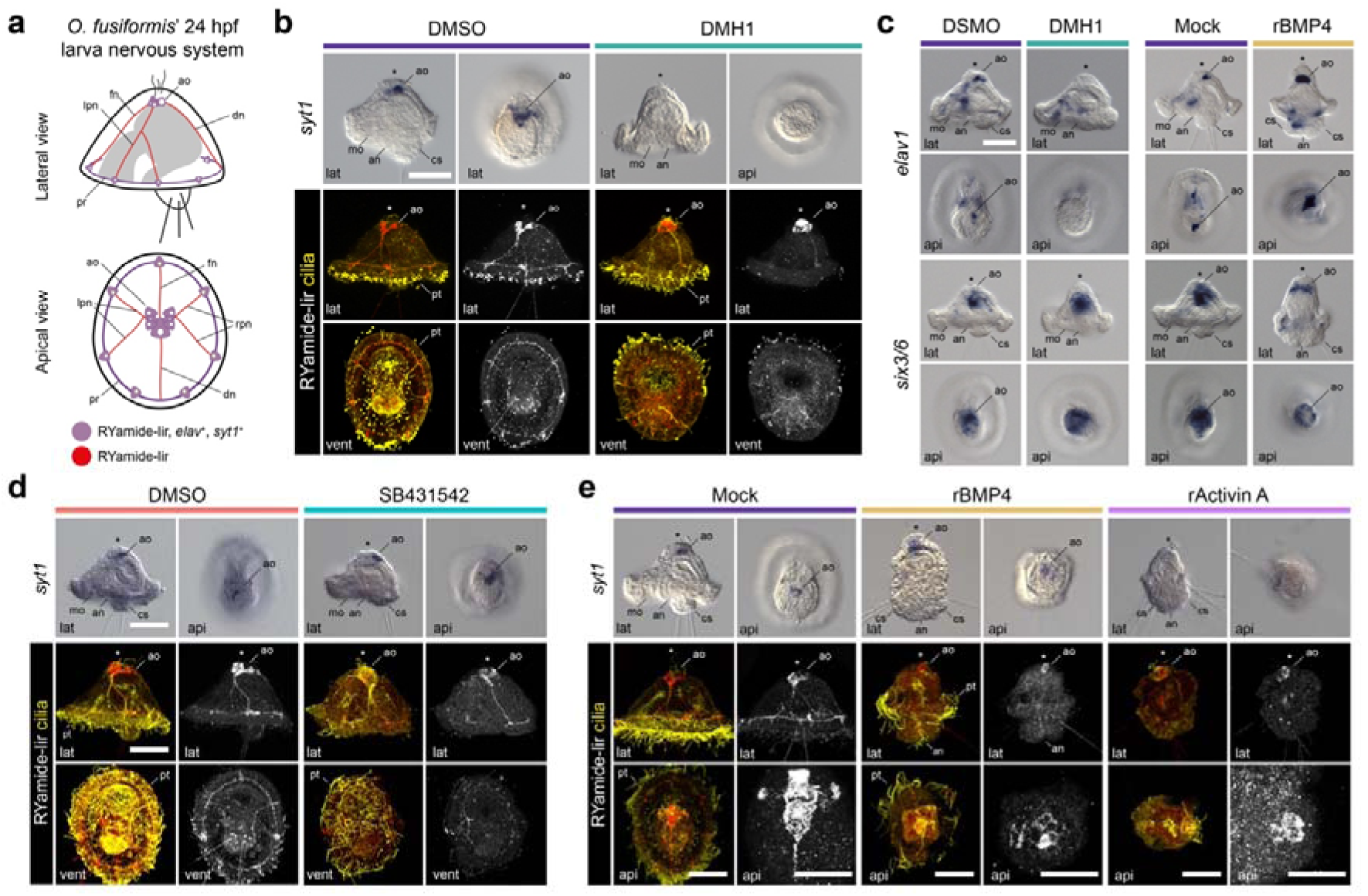
The role of BMP and Activin/Nodal in neural specification in *O. fusiformis*. (**a**) Schematic diagram of the larval nervous system at 24 hpf in *O. fusiformis*. (**b**) Whole mount *in situ* hybridisation of the neuronal marker *syt1* (top) and z-stack projections of RYamide immunoreactivity in control and treated larvae developed from embryos treated from 4 to 6 hpf with DMH1. (**c**) Whole mount *in situ* hybridisation of the neural markers *elav1* and *six3/6* in control and treated larvae developed from embryos treated from 4 to 6 hpf with DMH1 and rBMP4. (**d**–**e**) Whole mount *in situ* hybridisation of the neuronal marker *syt1* (top) and z-stack projections of RYamide immunoreactivity in control and treated larvae developed from embryos treated from 4 to 6 hpf with the Activin/Nodal inhibitor SB431542 (**d**) and overactivated with rBMP4 and rActivin A (**e**). In all panels, the asterisks mark the apical pole. Scale bars are 50 µm. an anus, ao apical organ, api apical, at apical tuft, ch chatae, cs chaetal sac, dn dorsal nerve, fn frontal nerve, lat lateral, lpn left peripheral nerve, mg midgut, mo mouth, pr prototrochal ring, pt prototroch, rpn right peripheral nerve, vent ventral. Drawings are not to scale.

**Extended Data Fig 4.**
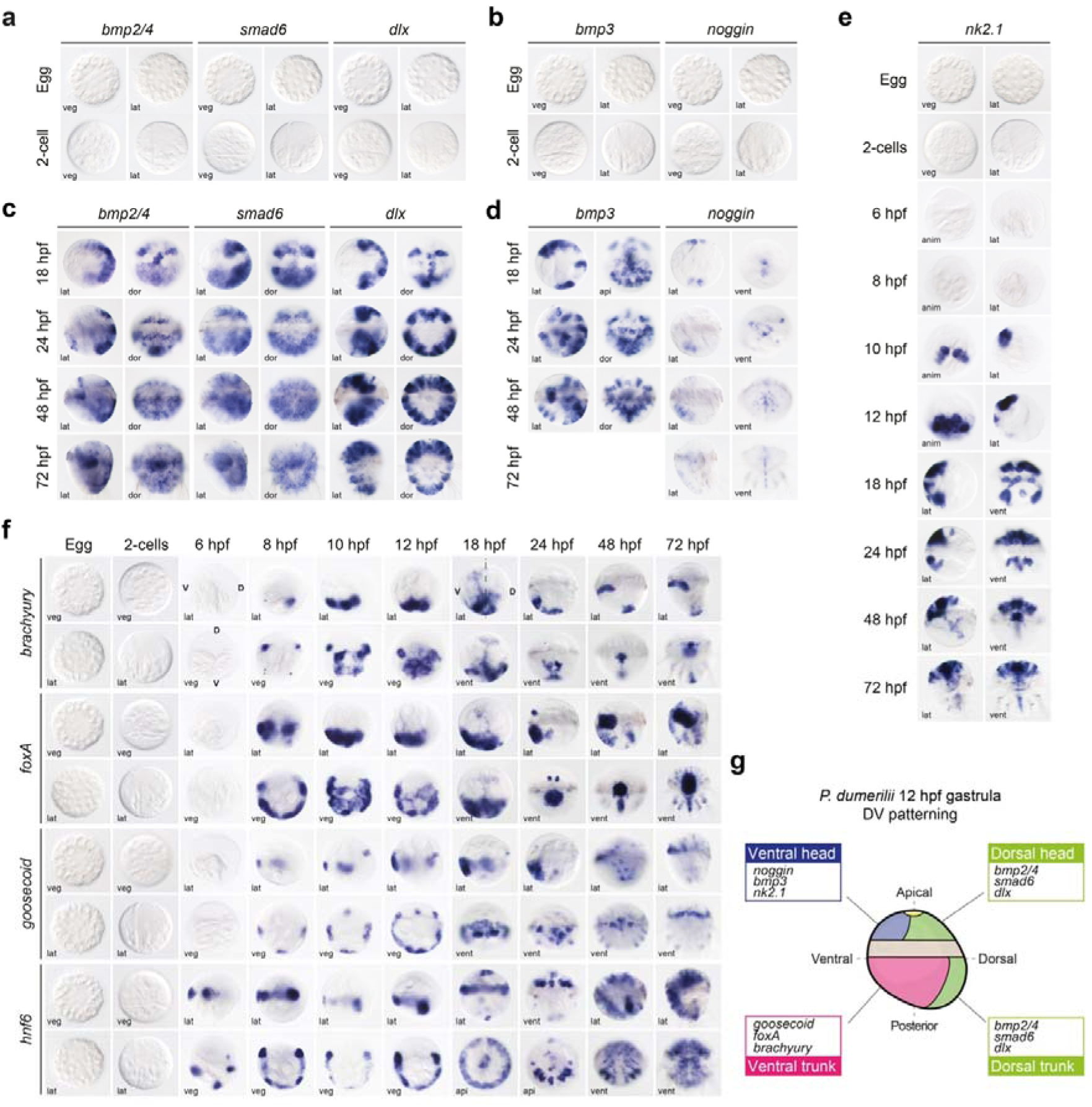
Expression of dorsoventral marker genes in *P. dumerilii.* (**a**–**e**) Whole mount *in situ* hybridisation of dorsoventral markers at selected embryonic and larval stages. (**f**) Whole mount *in situ* hybridisation of *brachyury*, *foxA*, *goosecoid*, and *hnf6* at selected embryonic and larval stages. (**g**) Diagram of dorsoventral marker genes, and *goosecoid*, *foxA*, and *brachyury* expression domains in a 12 hpf gastrula stage. A ciliary ring (the prototroch; grey) separates the anterior head and the posterior trunk. anim animal, veg vegetal, vent ventral, lat lateral. Drawing is not to scale.

**Extended Data Fig. 5.**
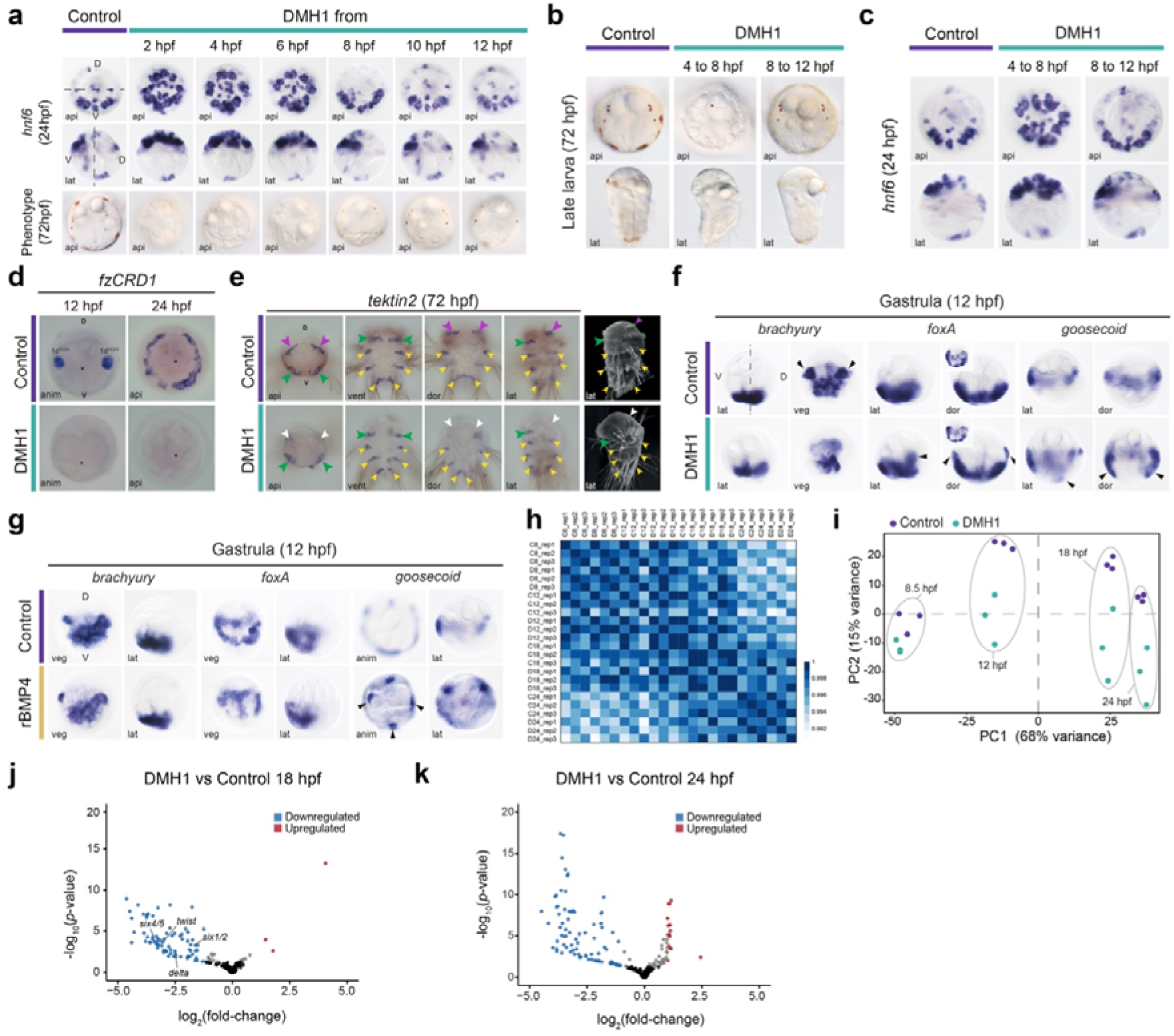
The role of BMP in dorsoventral axis specification in *Platynereis*. (**a**) Whole mount *in situ* hybridisation of *hnf6* at 24 hpf (two top row) and Differential Interference Contrast (DIC) images at 72 hpf (bottom row) in controls (first column) and different windows of treatments with DMH1. (**b**–**c**) DIC images (**b**) and whole mount *in situ* hybridisation (**c**) of *hnf6* in control and DMH1-treated in windows that result in an abnormal dorsoventral axis (4 to 8 hpf treatment) and wild-type phenotype (8 to 12 hpf treatment). (**d**) Whole mount *in situ* hybridisation of nephroblast (1c^11221^& 1d^11221^)/headkidney marker *fzCRD1* in control and DMH1-treated 12 and 24 hpf larvae. (**e**) Whole mount *in situ* hybridisation of the ciliary marker *tektin-2* and scanning electron microscopy images of control and DMH1-treated 72 hpf larvae. Ciliary bands like the dorsoanterior akrotroch (present/red, absent/white), the metatroch (green), and the paratrochs (yellow) are indicated by arrows. (**f**–**g**) Whole mount *in situ* hybridisation of *brachyury*, *foxA* and *goosecoid* in control and DMH1 (**f**) and rBMP4-treated (**g**) gastrula. (**h**) Hierarchically clustered pairwise correlation matrix between DMH1-treated and control samples. The scale shows the relative difference in Euclidean distance. (**i**) Principal component (PC) analysis plot for DMH1- treated vs control RNA-seq analyses. (**j**–**k**) Volcano plots depicting differentially expressed genes in 18 and 24 hpf larvae after DMH1 treatment from 4 to 8 hpf. anim animal, api apical, dor dorsal, lat lateral, veg vegetal.

**Extended Data Fig. 6.**
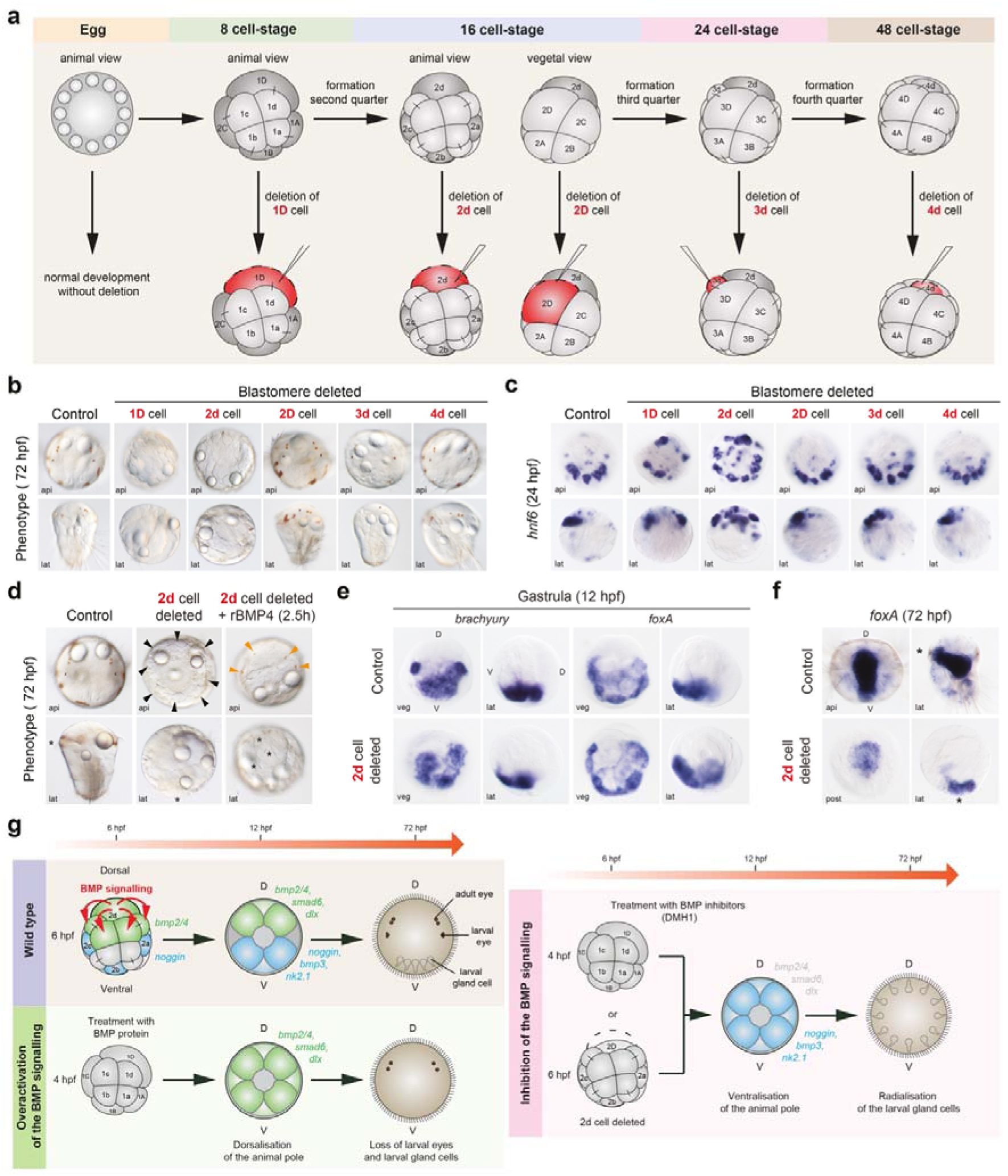
Effect of D-lineage blastomere ablations on dorsoventral specification in *Platynereis*. (**a**) Diagram of the blastomere ablations (red) at different cleavage stages. Ablated cells (1D is the precursor cell of 2d and 2D; 2D will bud off 3d and 4d) are indicated in red. (**b**) Differential Interference Contrast (DIC) images of 72 hpf larvae resulting from the blastomere ablations. (**c**) Whole mount *in situ* hybridisation of *hnf6* in larvae (24 hpf) after blastomere ablations. (**d**) DIC images of larvae rescued with rBMP4 after the 2d-cell ablation. At 72 hpf, larvae after 2d cell deletion exhibit the radialisation of gland cells (black arrowheads) and loss of eyes. Rescue with rBMP4 results in a reduction of gland cells and the retention of two pairs of adult eyes (orange arrowheads). (**e**) Whole mount *in situ* hybridisation of *brachyury* and *foxA* in gastrulae (12 hpf) after 2d ablation. (**f**) Whole mount *in situ* hybridisation of *foxA* in larvae (72 hpf) after 2d ablation. (**g**) Diagram of the role of BMP signalling in wild type, rBMP4- and DMH1-treated, and 2d cell-ablated embryos, depicting the effects on dorsoventral marker gene expression at the gastrula stage (12 hpf), and the effects on morphology at 72 hpf larva. The asterisks mark the mouth. api apical, D dorsal, lat lateral, post posterior, V ventral, veg vegetal.

**Extended Data Fig. 7.**
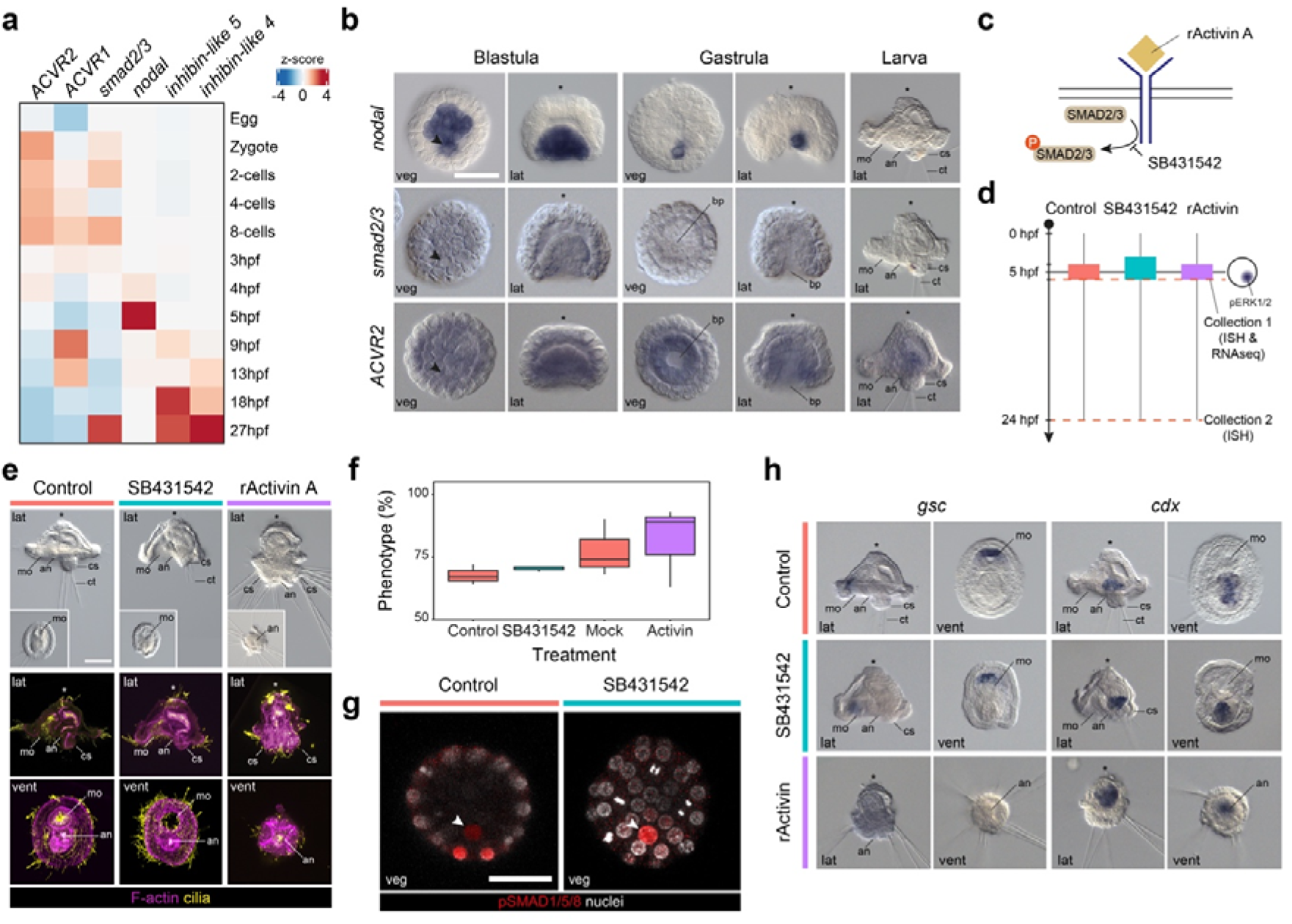
Activin/Nodal plays a role in dorsoposterior specification in *O. fusiformis* independently of pSMAD1/5/8. (**a**) Heatmap indicating the relative expression of core components of the Activin/Nodal pathway during the development of *O. fusiformis*. (**b**) Whole mount *in situ* hybridisation of the ligand *nodal*, secondary messenger *smad2/*3, and the activin receptor *ACVR2* in blastula (6 hpf), gastrula (9 hpf), and larval stages (24 hpf). (**c**) Simplified schematic of the Activin/Nodal pathway and mode of action of SB431542 and recombinant Activin A (rActivin A). (**d**) Schematic representation of the experimental design for the treatment windows and sample collections. (**e**) Differential interference contrast (top) and z-stack projections of control and treated larvae developed from embryos treated with SB431542 (3 to 6 hpf) and rActivin A (4 to 6 hpf). Insets in the first row are ventral views. Cilia (yellow) are labelled with tubulin, and F-actin (magenta) is labelled with phalloidin. (**f**) Box plots indicating the percentage of embryos exhibiting the described phenotypes for each treatment. DMSO and mock control larvae have a wild-type morphology. SB431542-treated larvae have a mild reduction of the dorsoposterior region, and rActivin-treated larvae have ectopic dorsoposterior tissue. (**g**) Z-stack projections of blastula (6 hpf) treated with control and SB431542/rActivin A stained against pSMAD1/5/8. (**h**) Whole mount *in situ* hybridisation of anterior (*gsc*) and posterior (*cdx*) marker genes in control and treated larvae developed from SB431542 and rActivin A-treated embryos. In all panels, the arrowheads point to the 4d organiser, and the asterisks mark the animal/apical pole. Scale bars are 50 µm. an anus, bp blastopore, ch chatae, cs chaetal sac, lat lateral, mo mouth, pt prototroch, veg vegetal, vent ventral.

**Extended Data Fig. 8.**
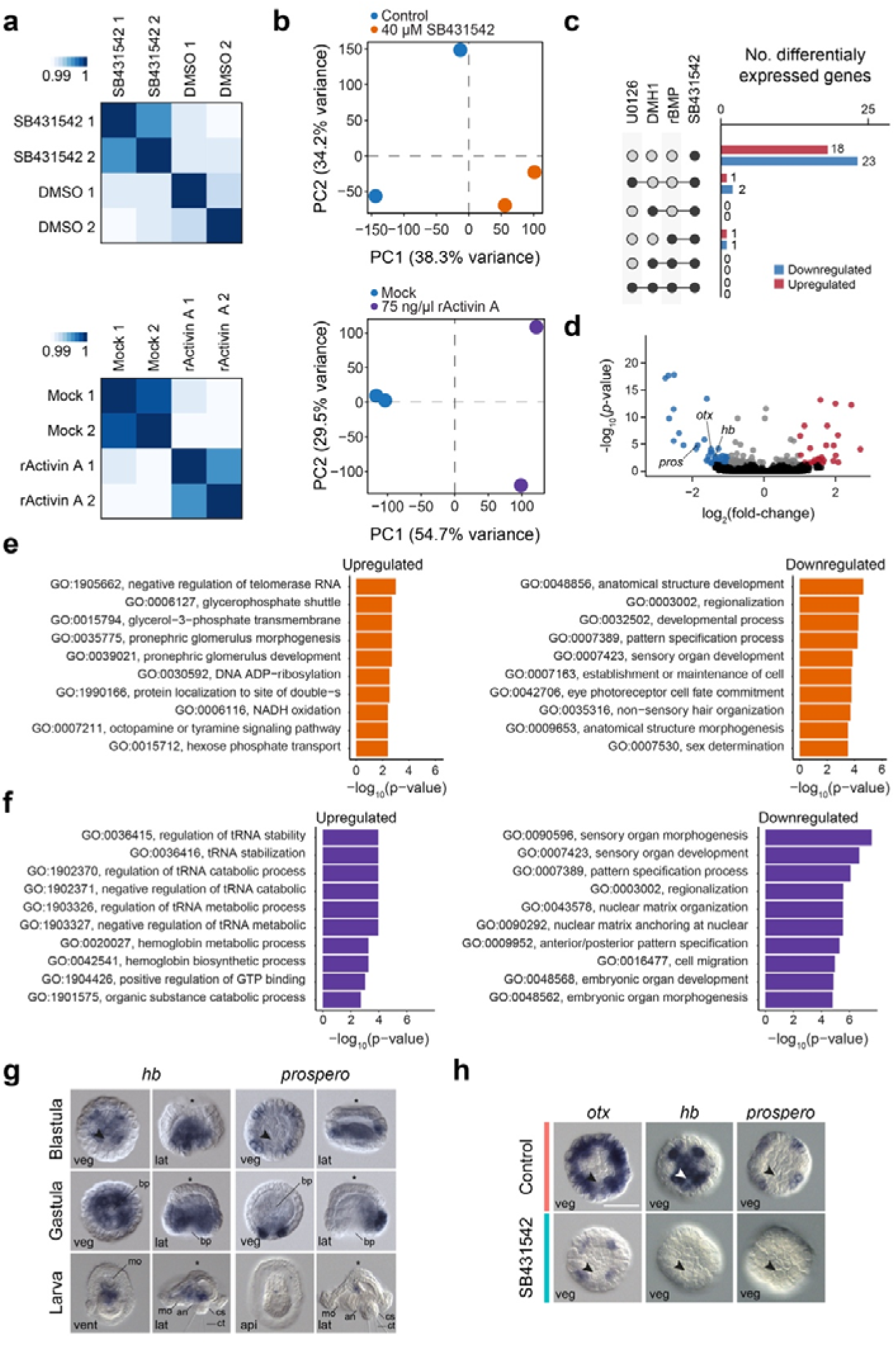
The gene regulatory network downstream of Activin/Nodal pathway in *O. fusiformis*. (**a**) Hierarchically clustered pairwise correlation matrix between SB431542 (top) and rActivin A (down) treated and control samples. The scale shows the relative difference in Euclidean distance. (**b**) Principal component (PC) analysis plots for SB431542 (top) and rActivin (bottom) transcriptomic analyses indicate that the main source of transcriptional variability between samples is the treatment condition (PC1). (**c**) Bar plots indicating the number of differentially expressed genes in different treatments. Full circles connected with a line indicated shared differentially expressed genes. (**d**) Volcano plot depicting differentially expressed genes after SB431542 treatment in 6 hpf blastulae. (**e**, **f**) Bar plots showing the top ten enriched Gene Ontology terms in differentially expressed genes after SB431542 (**e**) and rActivin A treatment (**f**). (**g**, **h**) Whole mount *in situ* hybridisation of selected candidate differentially expressed genes in SB431542-treated embryos at the blastula, gastrula, and larval stages. In (**g**, **h**), the arrowheads point to the 4d organiser, and the asterisks mark the animal/apical pole. Scale bars are 50 µm. an anus, bp blastopore, ch chatae, cs chaetal sac, lat lateral, mo mouth, pt prototroch, veg vegetal.

**Extended Data Fig. 9.**
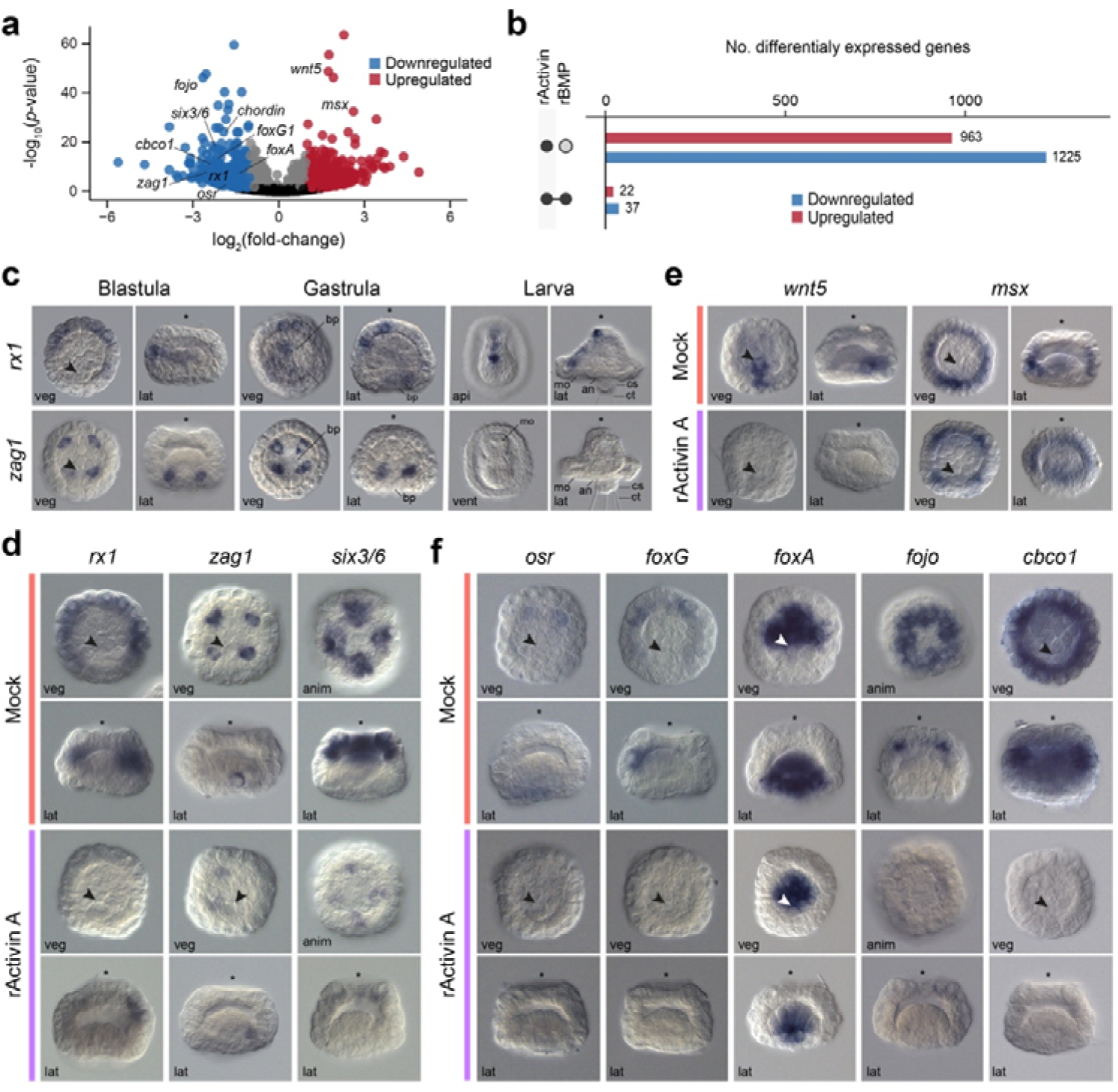
Differential gene expression analyses in recombinant Activin A- treated *O. fusiformis* embryos. (**a**) Volcano plot depicting differentially expressed genes after rActivin A treatment in 6 hpf blastulae. (**b**) Bar plots indicating the number of differentially expressed genes in different treatments. Full circles connected with a line indicated shared differentially expressed genes. (**c**) Whole mount *in situ* hybridisation of selected candidate differentially expressed genes after rActivin A treatment in wild type embryos at blastula, gastrula, and larval stages. (**e**–**f**) Whole mount *in situ* hybridisation of candidate genes downregulated only in recombinant Activin A-treated blastula (**e**), upregulated in rBMP4 and rActivin A-treated blastula (**f**), and downregulated in rBMP4 and rActivin A-treated blastula. In (**b**–**f**), the arrowheads point to the 4d organiser, and the asterisks mark the animal/apical pole. Scale bars are 50 µm. an anus, anim animal, bp blastopore, ch chatae, cs chaetal sac, lat lateral, mo mouth, veg vegetal.

**Extended Data Fig. 10.**
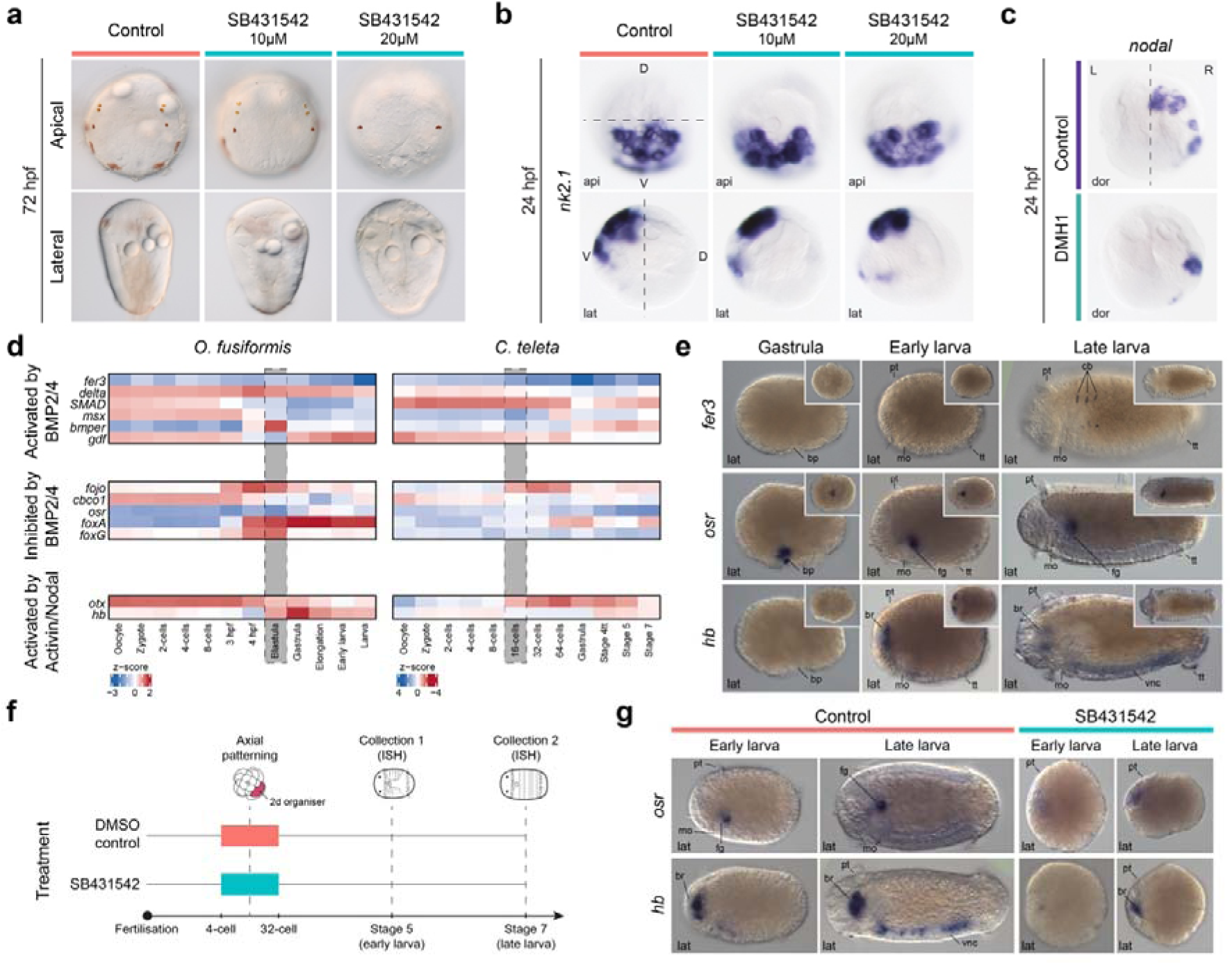
Activin/Nodal does not control dorsoventral polarity in *P. dumerilii* and uses a different downstream network in *C. teleta*. (**a**) Differential interference contrast images of control and SB431542 larvae at 72 hpf in *P. dumerilii*. (**b**) Whole mount *in situ* hybridisation of the ventral marker *nk2.1* in control and SB431542 24 hpf larvae of *P. dumerilii*. The dashed line marks the boundary between the dorsal (D) and ventral (V) sides. (**c**) Whole mount *in situ* hybridisation of *nodal* in control and DMH1- treated 24 hpf larvae in *P. dumerilii*. The dashed line marks the boundary between the left (L) and right (R) sides. (**d**) Heatmaps depicting the relative expression of differentially expressed candidate genes in *O. fusiformis* and their corresponding one-to-one orthologs in *C. teleta* during the embryogenesis of these species. (**e**) Whole mount *in situ* hybridisation of three candidate differentially expressed genes in *O. fusiformis* during the development of *C. teleta* at the gastrula, stage 5 and stage 7 larvae. Insets are ventral views. (**f**) Schematic diagram of the drug treatment in *C. teleta*. (**g**) Whole mount *in situ* hybridisation of the candidate genes *osr* (inhibited by the BMP pathway) and *hb* (activated by Activin/Nodal) in control and SB431542 stage 5 and stage 7 *C. teleta* larvae. Scale bars are 50 µm. an anus, api apical, bp blastopore, br brain, cb chaetoblast, dor dorsal, fg foregut, lat lateral, mo mouth, pt prototroch, tt telotroch, vnc ventral nerve cord.

## Supplementary Tables

**Supplementary Table 1.** Scoring of pSMAD1/5/8 immunoreactivity in blastula (6 hpf) of *O. fusiformis* embryos after U0126, DMH1, SB-431542, rBMP4 and rActivin A treatments.

**Supplementary Table 2.** Scoring of *chordin* and *bmp2/4* expression in larvae of *O. fusiformis* after U0126 treatment.

**Supplementary Table 3.** Scoring of morphological phenotypes in larvae of *O. fusiformis* after DMH1, SB431542, rBMP4 and rActivin A treatments.

**Supplementary Table 4.** Scoring of morphological phenotypes in *O. fusiformis* larvae after different treatment windows with DMH1, SB431542, rBMP4 and rActivin A.

**Supplementary Table 5.** Scoring of *cdx* and *gsc* expression in larvae of *O. fusiformis* larvae after different treatment windows with DMH1, SB431542, rBMP4 and rActivin A.

**Supplementary Table 6.** Scoring of *notch-like* and *BAMBI* expression in *O. fusiformis* larvae after DMH1, SB-431542, rBMP4 and rActivin A treatment.

**Supplementary Table 7.** Number of differentially expressed genes in the blastula (6 hpf) of *O. fusiformis* after DMH1, SB431542, rBMP4 and rActivin A treatments.

**Supplementary Table 8.** Differentially upregulated genes in the blastula (6 hpf) of *O. fusiformis* after DMH1 treatment.

**Supplementary Table 9.** Differentially downregulated genes in the blastula (6 hpf) of *O. fusiformis* after DMH1 treatment.

**Supplementary Table 10.** Differentially upregulated genes in the blastula (6 hpf) of *O. fusiformis* after rBMP4 treatment.

**Supplementary Table 11.** Differentially downregulated genes in the blastula (6 hpf) of *O. fusiformis* after rBMP4 treatment.

**Supplementary Table 12.** Scoring of the expression of candidate genes at the blastula stage of *O. fusiformis* after DMH1 treatment.

**Supplementary Table 13.** Scoring of the expression of candidate genes at the blastula stage of *O. fusiformis* after rBMP4 and rActivin A treatment.

**Supplementary Table 14.** Scoring of the expression of candidate genes at the blastula stage of *O. fusiformis* after SB431542 treatment.

**Supplementary Table 15.** Scoring of *syt1*, *elav1* and *six3/6* expression in larvae of *O. fusiformis* after DMH1, SB431542, rBMP4 and rActivin A treatment.

**Supplementary Table 16.** Scoring of RYamide-like immunoreactivity in larvae of *O. fusiformis* after DMH1, SB431542, rBMP4 and rActivin A treatment.

**Supplementary Table 17.** Scoring of dp-ERK1/2 immunoreactivity in blastulae of *S. lamarcki* treated with U0126 and DMH1.

**Supplementary Table 18.** Scoring of pSMAD1/5/8 immunoreactivity in blastulae of *S. lamarcki* treated with U0126 and DMH1.

**Supplementary Table 19.** Differentially downregulated genes in 8.5 hpf embryos of *P. dumerilii* after DMH1 treatment.

**Supplementary Table 20.** Differentially upregulated genes in 8.5 hpf embryos of *P. dumerilii* after DMH1 treatment.

**Supplementary Table 21.** Differentially downregulated genes in 12 hpf embryos of *P. dumerilii* after DMH1 treatment.

**Supplementary Table 22.** Differentially upregulated genes in 12 hpf embryos of *P. dumerilii* after DMH1 treatment.

**Supplementary Table 23.** Differentially downregulated genes in 18 hpf embryos of *P. dumerilii* after DMH1 treatment.

**Supplementary Table 24.** Differentially upregulated genes in 18 hpf embryos of *P. dumerilii* after DMH1 treatment.

**Supplementary Table 25.** Differentially downregulated genes in 24 hpf embryos of *P. dumerilii* after DMH1 treatment.

**Supplementary Table 26.** Differentially upregulated genes in 24 hpf embryos of *P. dumerilii* after DMH1 treatment.

**Supplementary Table 27.** Differentially upregulated genes in the blastula (6 hpf) of *O. fusiformis* after SB431542 treatment.

**Supplementary Table 28.** Differentially downregulated genes in the blastula (6 hpf) of *O. fusiformis* after SB431542 treatment.

**Supplementary Table 29.** Differentially upregulated genes in the blastula (6 hpf) of *O. fusiformis* after rActivin A treatment.

**Supplementary Table 30.** Differentially downregulated genes in the blastula (6 hpf) of *O. fusiformis* after rActivin A treatment.

**Supplementary Table 31.** Scoring of morphological phenotypes of stage 5 and 7 larvae of *C. teleta* treated with SB431542.

## References

1 Finnerty, J. R. The origins of axial patterning in the metazoa: how old is bilateral symmetry? Int J Dev Biol 47, 523–529 (2003).

2 Genikhovich, G. & Technau, U. On the evolution of bilaterality. Development 144, 3392–3404, doi:10.1242/dev.141507 (2017).

3 De Robertis, E. M. & Kuroda, H. Dorsal-ventral patterning and neural induction in *Xenopus* embryos. Annual Review of Cell and Developmental Biology 20, 285–308 (2004).

4 von Ohlen, T. & Doe, C. Q. Convergence of dorsal, dpp, and egfr signaling pathways subdivides the *Drosophila* neuroectoderm into three dorsal-ventral columns. Dev Biol 224, 362–372, doi:10.1006/dbio.2000.9789 (2000).

5 Morsdorf, D., Knabl, P. & Genikhovich, G. Highly conserved and extremely evolvable: BMP signalling in secondary axis patterning of Cnidaria and Bilateria. Dev Genes Evol 234, 1–19, doi:10.1007/s00427-024-00714-4 (2024).

6 Halanych, K. M. Evidence from 18S ribosomal DNA that the Lophophorates are protostome animal. Science 267, 1641–1643 (1995).

7 Seudre, O., Carrillo-Baltodano, A. M., Liang, Y. & Martín-Durán, J. M. ERK1/2 is an ancestral organising signal in spiral cleavage. Nat Commun 13, 2286, doi:10.1038/s41467-022-30004-4 (2022).

8 Lambert, J. D. Mesoderm in spiralians: the organizer and the 4d cell. J Exp Zool B Mol Dev Evol 310, 15–23, doi:10.1002/jez.b.21176 (2008).

9 Lambert, J. D., Johnson, A. B., Hudson, C. N. & Chan, A. Dpp/BMP2-4 mediates signaling from the D-quadrant organizer in a spiralian embryo. Curr Biol 26, 2003–2010, doi:10.1016/j.cub.2016.05.059 (2016).

10 Genikhovich, G. et al. Axis patterning by BMPs: Cnidarian network reveals evolutionary constraints. Cell Rep 10, 1646–1654, doi:10.1016/j.celrep.2015.02.035 (2015).

11 Seaver, E. C. Variation in spiralian development: insights from polychaetes. Int J Dev Biol 58, 457–467, doi:10.1387/ijdb.140154es (2014).

12 Tan, S., Huan, P. & Liu, B. Molluskan dorsal-ventral patterning relying on BMP2/4 and Chordin provides Insights into spiralian development and evolution. Mol Biol Evol 39, doi:10.1093/molbev/msab322 (2022).

13 Lyons, D. C., Perry, K. J., Batzel, G. & Henry, J. Q. BMP signaling plays a role in anterior-neural/head development, but not organizer activity, in the gastropod *Crepidula fornicata*. Dev Biol 463, 135–157, doi:10.1016/j.ydbio.2020.04.008 (2020).

14 Denes, A. S. et al. Molecular architecture of annelid nerve cord supports common origin of nervous system centralization in bilateria. Cell 129, 277–288, doi:10.1016/j.cell.2007.02.040 (2007).

15 Lanza, A. R. & Seaver, E. C. Functional evidence that Activin/Nodal signaling is required for establishing the dorsal-ventral axis in the annelid *Capitella teleta*. Development 147, doi:10.1242/dev.189373 (2020).

16 Lanza, A. R. & Seaver, E. C. An organizing role for the TGF-beta signaling pathway in axes formation of the annelid *Capitella teleta*. Dev Biol 435, 26–40, doi:10.1016/j.ydbio.2018.01.004 (2018).

17 Kuo, D. H. & Weisblat, D. A. A new molecular logic for BMP-mediated dorsoventral patterning in the leech *Helobdella*. Curr Biol 21, 1282–1288, doi:10.1016/j.cub.2011.06.024 (2011).

18 Martín-Zamora, F. M. et al. Annelid functional genomics reveal the origins of bilaterian life cycles. Nature 615, 105–110, doi:10.1038/s41586-022-05636-7 (2023).

19 Lanza, A. R. & Seaver, E. C. Activin/Nodal signaling mediates dorsal-ventral axis formation before third quartet formation in embryos of the annelid *Chaetopterus pergamentaceus*. Evodevo 11, 17, doi:10.1186/s13227-020-00161-y (2020).

20 Martin-Duran, J. M. & Marletaz, F. Unravelling spiral cleavage. Development 147, doi:10.1242/dev.181081 (2020).

21 Andrikou, C. & Hejnol, A. FGF signaling acts on different levels of mesoderm development within Spiralia. Development 148, doi:10.1242/dev.196089 (2021).

22 Lewin, T. D. et al. Brachiopod genome unveils the evolution of the BMP–Chordin network in bilaterian body patterning. BiorXiv, doi:10.1101/2024.05.28.596352 (2024).

23 Martín-Durán, J. M., Passamaneck, Y. J., Martindale, M. Q. & Hejnol, A. The developmental basis for the recurrent evolution of deuterostomy and protostomy. Nat Ecol Evol 1, 5, doi:10.1038/s41559-016-0005 (2016).

24 Tan, S., Huan, P. & Liu, B. Functional evidence that FGFR regulates MAPK signaling in organizer specification in the gastropod mollusk *Lottia peitaihoensis*. Mar Life Sci Technol 5, 455–466, doi:10.1007/s42995-023-00194-x (2023).

25 Moggioli, G. et al. Distinct genomic routes underlie transitions to specialised symbiotic lifestyles in deep-sea annelid worms. Nat Commun 14, 2814, doi:10.1038/s41467-023-38521-6 (2023).

26 Mörsdorf, D., Prünster, M. M. & Genikhovich, G. Chordin-mediated BMP shuttling patterns the secondary body axis in a cnidarian. BiorXiv, doi:10.1101/2024.05.27.596067 (2024).

27 Kretzschmar, M., Doody, J. & Massagu, J. Opposing BMP and EGF signaling pathways converge ono the TGF-b family mediator Smad1. Nature 389, 618–622 (1997).

28 Sapkota, G., Alarcon, C., Spagnoli, F. M., Brivanlou, A. H. & Massague, J. Balancing BMP signaling through integrated inputs into the Smad1 linker. Mol Cell 25, 441–454, doi:10.1016/j.molcel.2007.01.006 (2007).

29 Fuentealba, L. C. et al. Integrating patterning signals: Wnt/GSK3 regulates the duration of the BMP/Smad1 signal. Cell 131, 980–993, doi:10.1016/j.cell.2007.09.027 (2007).

30 Carrillo-Baltodano, A. M. & Meyer, N. P. Decoupling brain from nerve cord development in the annelid *Capitella teleta*: Insights into the evolution of nervous systems. Dev Biol 431, 134–144, doi:10.1016/j.ydbio.2017.09.022 (2017).

31 Carrillo-Baltodano, A. M., Donnellan, R. D., Williams, E. A., Jekely, G. & Martin-Duran, J. M. The development of the adult nervous system in the annelid *Owenia fusiformis*. Neural Dev 19, 3, doi:10.1186/s13064-024-00180-8 (2024).

32 Carrillo-Baltodano, A. M., Seudre, O., Guynes, K. & Martín-Durán, J. M. Early embryogenesis and organogenesis in the annelid *Owenia fusiformis*. Evodevo 12, 5, doi:10.1186/s13227-021-00176-z (2021).

33 Conzelmann, M. & Jékely, G. Antibodies against conserved amidated neuropeptide epitopes enrich the comparative neurobiology toolbox. Evodevo 3, 1–11 (2012).

34 Conzelmann, M. et al. Neuropeptides regulate swimming depth of *Platynereis* larvae. Proc Natl Acad Sci U S A 108, E1174–1183, doi:10.1073/pnas.1109085108 (2011).

35 Williams, E. A. et al. Synaptic and peptidergic connectome of a neurosecretory center in the annelid brain. Elife 6, doi:10.7554/eLife.26349 (2017).

36 Helm, C., Vocking, O., Kourtesis, I. & Hausen, H. *Owenia fusiformis* - a basally branching annelid suitable for studying ancestral features of annelid neural development. BMC Evol Biol 16, 129, doi:10.1186/s12862-016-0690-4 (2016).

37 Namigai, E. K. O. & Shimeld, S. M. Live imaging of cleavage variability and vesicle flow dynamics in dextral and sinistral spiralian embryos. Zoolog Sci 36, 5–16, doi:10.2108/zs180088 (2019).

38 Lambert, J. D. & Nagy, L. M. The MAPK cascade in equally cleaving spiralian embryos. Dev Biol 263, doi:10.1016/S0012-1606(03)00438-X (2003).

39 McDougall, C., Chen, W. C., Shimeld, S. M. & Ferrier, D. E. The development of the larval nervous system, musculature and ciliary bands of *Pomatoceros lamarckii* (Annelida): heterochrony in polychaetes. Front Zool 3, 16, doi:10.1186/1742-9994-3-16 (2006).

40 Özpolat, B. D. et al. The Nereid on the rise: *Platynereis* as a model system. Evodevo 12, 10, doi:10.1186/s13227-021-00180-3 (2021).

41 Wilson, E. B. The cell-lineage of *Nereis*. A contribution to the cytogeny of the annelid body. Journal of Morphology 6, 361–480 (1892).

42 Dorresteijn, A. Quantitative analysis of cellular differentiation during early embryogenesis of *Platynereis dumerilii*. Roux’s Archiv Dev Biol 199, 14–30 (1990).

43 Vopalensky, P., Tosches, M. A., Achim, K., Handberg-Thorsager, M. & Arendt, D. From spiral cleavage to bilateral symmetry: the developmental cell lineage of the annelid brain. BMC Biol 17, 81, doi:10.1186/s12915-019-0705-x (2019).

44 Pfeifer, K., Schaub, C., Domsch, K., Dorresteijn, A. & Wolfstetter, G. Maternal inheritance of *twist* and analysis of MAPK activation in embryos of the polychaete annelid *Platynereis dumerilii*. PLoS One 9, e96702, doi:10.1371/journal.pone.0096702 (2014).

45 Luo, T., Matsuo-Takasaki, M., Lim, J. H. & Sargent, T. D. Differential regulation of *Dlx* gene expression by a BMP morphogenetic gradient. Int J Dev Biol 45, 681–684 (2001).

46 Bastin, B. R., Meha, S. M., Khindurangala, L. & Schneider, S. Q. Cooption of regulatory modules for tektin paralogs during ciliary band formation in a marine annelid larva. Dev Biol 503, 95–110, doi:10.1016/j.ydbio.2023.07.006 (2023).

47 Bastin, B. R. & Schneider, S. Q. Taxon-specific expansion and loss of *tektins* inform metazoan ciliary diversity. BMC Evol Biol 19, 40, doi:10.1186/s12862-019-1360-0 (2019).

48 Webster, N. B. & Meyer, N. P. *Capitella teleta* gets left out: possible evolutionary shift causes loss of left tissues rather than increased neural tissue from dominant-negative BMPR1. Neural Dev 19, 4, doi:10.1186/s13064-024-00181-7 (2024).

49 De Robertis, E. M. & Tejeda-Munoz, N. Evo-Devo of Urbilateria and its larval forms. Dev Biol 487, 10–20, doi:10.1016/j.ydbio.2022.04.003 (2022).

50 Amiel, A. R., Henry, J. Q. & Seaver, E. C. An organizing activity is required for head patterning and cell fate specification in the polychaete annelid *Capitella teleta*: new insights into cell-cell signaling in Lophotrochozoa. Dev Biol 379, 107–122, doi:10.1016/j.ydbio.2013.04.011 (2013).

51 Boyle, M. J., Yamaguchi, E. & Seaver, E. C. Molecular conservatioin of metazoan gut formation: evidence from expression of endomesoderm genes in *Capitella teleta* (Annelida). EvoDevo 5, 1–19 (2014).

52 Liang, Y., Wei, J., Kang, Y., Carrillo-Baltodano, A. M. & Martín-Durán, J. M. Cell fate specification modes shape transcriptome evolution in the highly conserved spiral cleavage. BiorXiv, 1–48, doi:10.1101/2024.12.25.630330 (2024).

53 Webster, N. B., Corbet, M., Sur, A. & Meyer, N. P. Role of BMP signaling during early development of the annelid *Capitella teleta*. Dev Biol 478, 183–204, doi:10.1016/j.ydbio.2021.06.011 (2021).

54 McColgan, A. & DiFrisco, J. Understanding developmental system drift. Development 151, doi:10.1242/dev.203054 (2024).

55 True, J. R. & Haag, E. S. Developmental system drift and flexibility in evolutionary trajectories. Evol Dev 3, 109–119, doi:10.1046/j.1525-142x.2001.003002109.x (2001).

56 Floc’hlay, S. et al. Deciphering and modelling the TGF-beta signalling interplays specifying the dorsal-ventral axis of the sea urchin embryo. Development 148, doi:10.1242/dev.189944 (2021).

57 Lapraz, F., Besnardeau, L. & Lepage, T. Patterning of the dorsal-ventral axis in echinoderms: insights into the evolution of the BMP-chordin signaling network. PLoS Biol 7, e1000248, doi:10.1371/journal.pbio.1000248 (2009).

58 Hill, C. S. Establishment and interpretation of NODAL and BMP signaling gradients in early vertebrate development. Curr Top Dev Biol 149, 311–340, doi:10.1016/bs.ctdb.2021.12.002 (2022).

59 Lambert, J. D. & Nagy, L. M. Asymmetric inheritance of centrosomally localized mRNAs during embryonic cleavages. Nature 420, 682–686 (2002).

60 Meyer, N. P., Carrillo-Baltodano, A., Moore, R. E. & Seaver, E. C. Nervous system development in lecithotrophic larval and juvenile stages of the annelid *Capitella teleta*. Front Zool 12, 15, doi:10.1186/s12983-015-0108-y (2015).

61 Seaver, E. C. Growth patterns during segmentation in the two polychaete annelids, Capitella sp. I and Hydroides elegans comparisons at distinct. (2005).

62 Chen, S., Zhou, Y., Chen, Y. & Gu, J. fastp: an ultra-fast all-in-one FASTQ preprocessor. Bioinformatics 34, i884–i890, doi:10.1093/bioinformatics/bty560 (2018).

63 Bray, N. L., Pimentel, H., Melsted, P. & Pachter, L. Near-optimal probabilistic RNA-seq quantification. Nat Biotechnol 34, 525–527, doi:10.1038/nbt.3519 (2016).

64 Love, M. I., Huber, W. & Anders, S. Moderated estimation of fold change and dispersion for RNA-seq data with DESeq2. Genome Biol 15, 550, doi:10.1186/s13059-014-0550-8 (2014).

65 Alexa, A. & Rahnenfuhrer, J. topGO: Enrichment analysis for gene ontology, <https://bioconductor.org/packages/release/bioc/html/topGO.html> (2024).

66 Bolger, A. M., Lohse, M. & Usadel, B. Trimmomatic: a flexible trimmer for Illumina sequence data. Bioinformatics 30, 2114–2120, doi:10.1093/bioinformatics/btu170 (2014).

67 Dobin, A. et al. STAR: ultrafast universal RNA-seq aligner. Bioinformatics 29, 15–21, doi:10.1093/bioinformatics/bts635 (2013).

68 Liao, Y., Smyth, G. K. & Shi, W. featureCounts: an efficient general purpose program for assigning sequence reads to genomic features. Bioinformatics 30, 923–930, doi:10.1093/bioinformatics/btt656 (2014).

